# Aggregated gene co-expression networks for predicting transcription factor regulatory landscapes in a non-model plant species

**DOI:** 10.1101/2023.04.24.538042

**Authors:** Luis Orduña-Rubio, Antonio Santiago, David Navarro-Payá, Chen Zhang, Darren C. J. Wong, J. Tomás Matus

**Author notes:** Shared corresponding authorship.

## Abstract

Gene co-expression networks (GCNs) have not been extensively studied in non-model plants. However, the rapid accumulation of transcriptome datasets in these species represents an opportunity to explore underutilized network aggregation approaches that highlight robust co-expression interactions and improve functional connectivity. We applied and evaluated two different aggregation methods on public grapevine RNA-Seq datasets belonging to three different tissue conditions (leaf, berry and ‘all organs’). Our results show that co-occurrence-based aggregation generally yielded the best-performing networks. We applied GCNs to study several TF gene families, showing its capacity of detecting both already-described and novel regulatory relationships between R2R3-MYBs, bHLH/MYC and multiple secondary metabolism pathway reactions. Specifically, TF gene-and pathway-centered network analyses successfully ascertained the previously established role of *VviMYBPA1* in controlling the accumulation of proanthocyanidins while providing insights into its novel role as a regulator of *p*-coumaroyl-CoA biosynthesis as well as the shikimate and aromatic amino-acid pathways. This network was validated using DNA Affinity Purification Sequencing data, demonstrating that co-expression networks of transcriptional activators can serve as a proxy of gene regulatory networks. This study presents an open repository to reproduce networks and a GCN application within the Vitviz platform, a user-friendly tool for exploring co-expression relationships.

## INTRODUCTION

Advances in genomics have enabled the exploitation of gene co-expression networks (GCNs) and their diverse approaches to hypothesize roles of uncharacterized genes in different plant model species (Burks et al., 2022; Madhamshettiwar et al., 2012). In this context, these networks are typically constructed with transcriptomic expression datasets (microarray or RNA-seq) where nodes represent genes and edges indicate co-expression relationships between them.

Among the currently available approaches for generating GCNs, those implying supervised methods, usually based on machine learning algorithms, have been shown to perform better with small networks (i.e., with a reduced number of genes, between tens to hundreds) in obtaining regulatory relationships between genes (Maetschke et al., 2014). Nevertheless, supervised methods require previous gene-function knowledge to be used as a training dataset, and prediction accuracy is lower when inferring genome-scale networks (Liesecke et al., 2018), making them less advisable in non-model species.

Networks constructed with unsupervised methods, instead, do not require previous knowledge as they capture transcriptional associations by retaining strongly-correlated genes. These GCNs are based on the "*guilt-by-association*" principle whereby correlation in gene expression implies biological association (Wolfe et al., 2005). More importantly, when applied to genome-scale networks (e.g. often > 10,000 genes), unsupervised methods do not require either time-consuming training processes or a previous feature selection step to reduce the size of the inference problem (Maetschke et al., 2014). Unsupervised methods are hence well-suited for gene identification in non-model species, where a low percentage of genes have been characterized but a large number of transcriptomic datasets are available. In addition, a few studies carried out in model species (e.g., in mouse and yeast, tomato, maize and arabidopsis) have shown that aggregation of individually-and independently-generated GCNs (Liesecke et al., 2019; Verleyen et al., 2015) can improve network performance when using unsupervised methods.

Among the many emerging non-model crop species, grapevine (*Vitis vinifera* L.), represents an ideal case for the implementation of GCNs due to the vast amount of data generated from transcriptomic methods, especially high-throughput next-generation sequencing techniques (Savoi et al., 2022). In grapevine, microarray-based GCNs have been generated (D. C. Wong et al., 2013) and applied to predict the function of individual genes or gene families (Loyola et al., 2016; Amato et al., 2017; Wong & Matus, 2017; Sun et al., 2018). Recently, aggregation across multiple individual microarray-based GCNs showed promising results of improved gene function prediction metrics compared to various types of atypical non-aggregated ones (Wong, 2020). However, RNA-Seq datasets have not been exploited in grapevine when it comes to GCN generation, even though RNA-Seq presents major advantages over microarrays, such as RNA-Seq being unbiased (i.e., only a fraction of genes are present in a microarray chip), which provides a higher dynamic range and whole transcriptome assessment (Hitzemann et al., 2013). Although recent studies have explored the potential of RNA-Seq in generating grapevine discrete GCNs (Pilati et al., 2021; Vannozzi et al., 2018), network aggregation approaches have not been evaluated further. Furthermore, there are many ways network aggregation across multiple GCN can be performed (Liesecke et al., 2018). Despite the limited pool of aggregation methods tested thus far, the HRR ranking of raw PCC values appears to be a reliable unsupervised approach for RNA-seq datasets. Additionally, there is still room for improving GCN generation in grapevine, such as the inclusion of tissue/organ-dependency, i.e., separating the datasets according to their tissue precedence, in order to generate whole genome-scale tissue-specific networks. Condition dependent networks may indeed reveal novel tissue-specific relationships between genes (Fukushima et al., 2012). Also, although there are some popular online tools for exploring GCNs in model species (e.g., in Arabidopsis, Szklarczyk et al., 2019; and tomato, Zouine et al., 2017), such resources are lacking for many crop plants such as grapevine.

Despite grapevine’s economic importance, the proportion of characterized genes, including large transcription factors (TFs) families, is comparatively low for model species such as *Arabidopsis thaliana* or *Solanum lycopersicum*. Nevertheless, the significant amount of available transcriptomic datasets in grapes represents a potential advantage for identifying regulatory networks, i.e., defining the intricate repertoire of target genes and processes controlled by a transcription factor (TF). Among all different types of TFs present in a cell (e.g., developmental or stress response regulators), those activating metabolic pathways could easily be identified by using GCNs since they generally present high co-expression values with their targets, i.e., structural genes of the metabolic pathways they control (Hu et al., 2020).

The present study comprises a deep analysis of different network construction methods, involving aggregation and condition dependency. Leaf, fruit (berry), and tissue-independent GCNs were computed and used as tools for the functional prediction of transcription factors by generating and analyzing single gene-centered networks for different transcription factor families. We ascertained that the PCC-HRR method is the best and most reliable approach for generating GCNs using a large dataset of publicly available RNA-Seq experiments. The pipelines developed are publicly available (https://github.com/Tomsbiolab) and can be adapted to any species of interest with a sufficiently large number of expression datasets available. As a result, we venture an important contribution of our method in proposing roles of uncharacterized genes in non-model plants. This would represent an effective tool to characterize both novel regulators of gene expression and their targets. Notwithstanding the numerous known and novel regulatory networks highlighted here, we take the opportunity to validate the robustness of our approach by leveraging a combination of state-of-the-art DNA affinity purification sequencing (DAP-Seq), transcriptomics, meta-analysis of TF-overexpression experiments, and literature support. Finally, this study also presents a GCN application within the Vitviz platform (www.vitviz.tomsbiolab.com), a user-friendly online tool for accessing, exploring and downloading co-expression relationships in grapevine.

## MATERIALS AND METHODS

### Generation of aggregated and single genome co-expression networks

A total of 4,815 transcriptomic RNA-Seq runs (Illumina sequencing) from grapevine public experiments were downloaded from NCBI Sequence Read Archive (SRA) (https://www.ncbi.nlm.nih.gov/sra). Metadata was first automatically and then manually classified according to their tissue precedence. SRA studies were manually inspected and filtered to keep those whose tissue precedence were correctly annotated while also excluding SRA studies concerning microRNAs, small RNAs and non-coding RNAs. Runs were trimmed with fastp version 0.20.0 (Chen et al., 2018) and aligned against the PN40024 reference genome assembly (12X.v2) for *V. vinifera* (Canaguier et al., 2017) with STAR aligner version 2.7.3a (Dobin et al., 2013). Raw counts for each run were computed using FeatureCounts version 2.0.0 (Liao et al., 2014) and the VCost.v3_27.gff3 gene models (available at http://www.integrape.eu/), generating a raw count matrix per SRA run. The code used for downloading, trimming, aligning and counting of raw counts is publicly available at https://github.com/Tomsbiolab/raw_counts_generation. A general overview of the pipeline is available at (Sup. Figure 1A).

**Figure 1.**
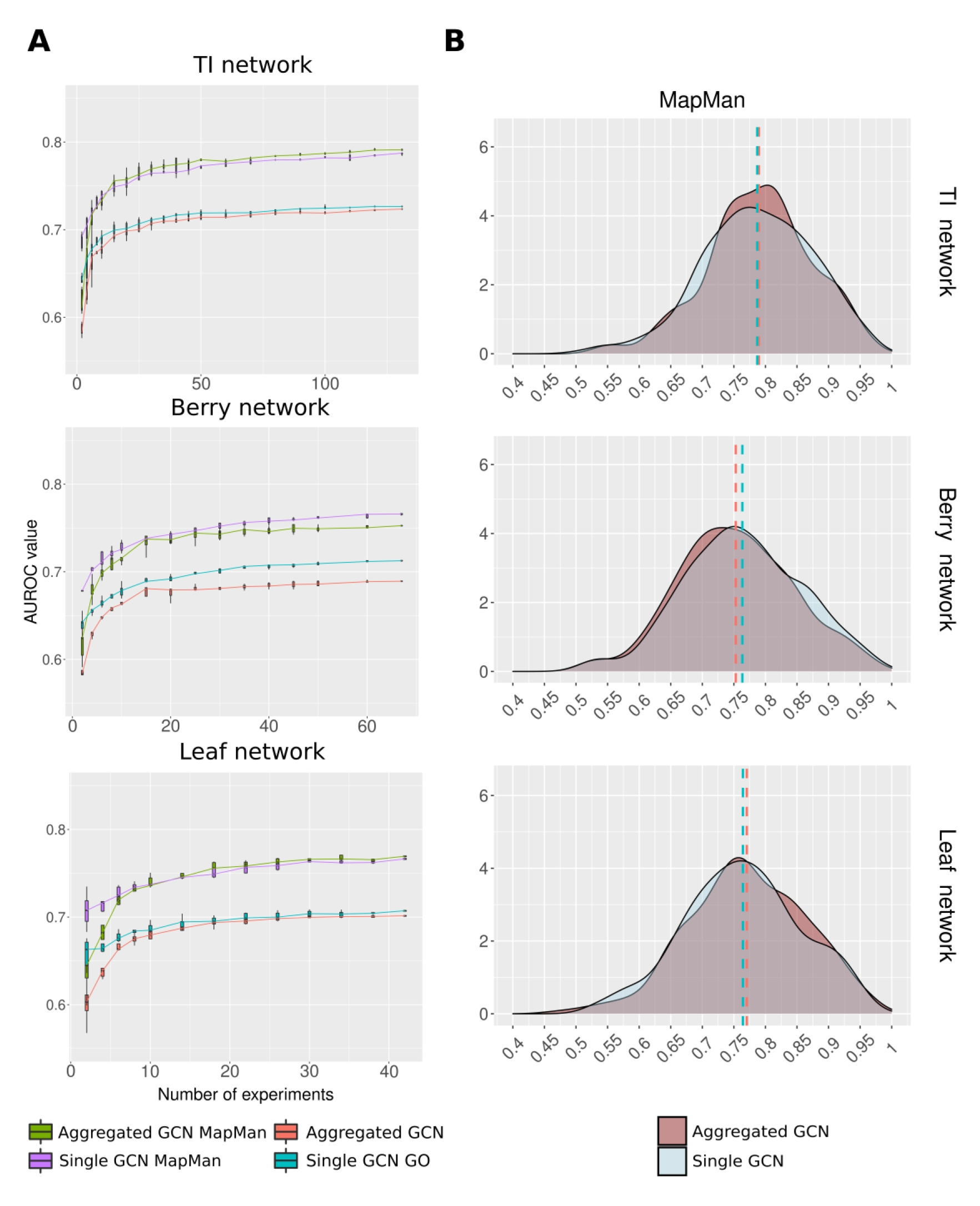
Co-occurrence-based aggregation generally improves network performance: A) Representation of AUROC values against the number of experiments across the berry, leaf and tissue-independent networks for all the single and aggregated GCNs with MapMan and GO ontologies. SRA study subsets were randomly selected in a cumulative manner, repeating the analysis five times (Sup. Figure 2). B) Density plot of the AUROC values for generated GCNs across each individual MapMan bin.

Aggregated GCN generation begins by selecting the SRA studies according to their tissue precedence. The pipeline then groups raw count matrices by SRA study, only considering runs with more than 10M alignments. SRA studies with less than 4 runs were discarded. In total, 131 SRA studies (2,766 runs) were used for the construction of the tissue-independent co-occurrence-based aggregation of GCNs (aggregated GCN), encompassing SRA studies from 9 different tissues (leaf, fruit, root, stem, wood, flower, seed, tendril and bud). 42 (670 runs) and 67 (1,615 runs) SRA studies were used for constructing the leaf-and berry-specific aggregated GCNs respectively. SRA studies were analyzed independently as described in (Orduña et al., 2022; D. C. J. Wong, 2020). Raw count matrices of the SRA studies were normalized to FPKM, and genes with less than 0.5 FPKMs in every run of the SRA study were discarded from the FPKM matrix. Pearson’s correlation coefficients of each gene against all other remaining genes were then calculated for each SRA study (across all individual SRA study FPKM matrices) and ranked in descending order. Ranked Pearson’s correlation coefficient (PCC) values were used to compute the highest reciprocal ranking (HRR) matrix (Mutwil et al., 2010), for each SRA study, considering only the top 420 ranked genes (420 roughly equals 1% of all VCost.v3 gene models), using the following formula: *HRR(A,B) = max(rank(A,B), rank(B,A))*. The aggregated GCN was then generated by computing the frequency of co-expression interaction(s) across individual HRR matrices. The frequency of co-expression was used as the edge weight in the GCN. Finally, as a noise-filtering step, only the top 420 frequency values for each gene were considered in building the final aggregate network, i.e., frequency values below this tier were changed to 0. In addition, the top 420 most highly co-expressed genes for any gene of interest were used to generate a gene-centered network of that particular gene. The code used for generating aggregated GCNs is available at https://github.com/Tomsbiolab/agg_WGCN. Similarly, the generation of the single network constructed from merged datasets, i.e., single GCNs, also followed the same pipeline with the only difference that all the SRA runs were grouped into a single raw count matrix. The code used for generating single GCNs is available at https://github.com/Tomsbiolab/non_agg_WGCN. A general overview of both GCN generation pipelines is available at (Sup. Figure 1B).

### Network performance evaluation

Network functional connectivity (i.e., performance) across all genes was assessed by neighbor-voting, a machine learning algorithm based on the guilt-by-association principle, which states that genes sharing common functions are often coordinately regulated across multiple experiments (Verleyen et al., 2015). The evaluation was performed using the EGAD R package (Ballouz et al., 2017) with default settings. The network was scored by the area under the receiver operating characteristics curve (AUROC) across updated MapMan (Thimm et al., 2004) V4 BIN functional categories and GO terms (http://geneontology.org/) associated with the grape genome reference VCost.v3 annotation using a threefold cross-validation. MapMan ontology generated in (Orduña et al., 2022) was edited by manually checking the genes present in the 9.2.2 bin and its sub-bins. MapMan BIN ontology annotations and GO terms were limited to groups containing 20–1000 genes to ensure robustness and stable performance when using the neighbor-voting algorithm, and MapMan BINs related to secondary metabolism were manually curated. Ontologies used for AUROC metric evaluation are available at Sup. Dataset 1.

### Evaluating the effect of network aggregation and the number of studies used

The impact of adding individual SRA networks/studies to aggregated and single GCNs respectively was evaluated by selecting random subsets of the used SRA studies in an additive manner and computing the AUROC of their resulting aggregated and single GCNs for each step. Importantly, the same exact subsets were selected for each type of network construction. The initial subset to build and evaluate GCNs consisted of two SRA studies after which further SRA study subsets were added, generating subsets of the following sizes: TI GCN (2, 5, and 10 increments for ranges 2 – 10, 15 – 50, 60 – 120, respectively), berry GCN (increments of 2 and 5 for ranges as above), and leaf GCN (2 and 4 increments for ranges 2 – 10 and 14 – 38, respectively). For each subset evaluated, this process was repeated 5 times. Additionally, the full set of SRA studies used for a particular GCN (i.e. 131, 67, and 42 for TI, berry, and leaf dataset, respectively) is also included. A general overview of the validation pipeline is available at (Sup. Figure 2)

**Figure 2.**
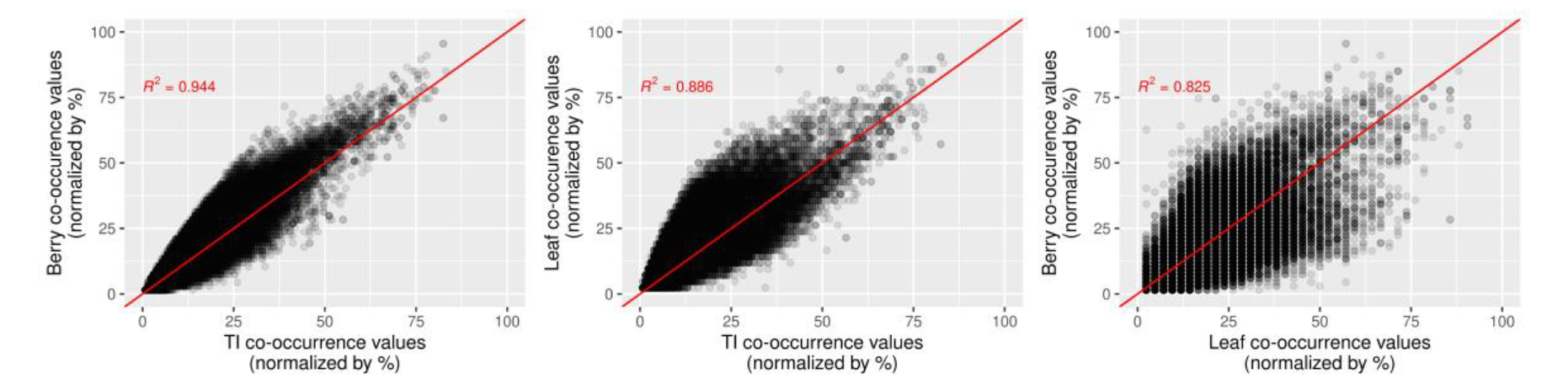
Berry and leaf GCNs differ more with respect to each other than when compared to the tissue-independent (TI) GCN. Normalized co-occurrence value comparisons by tissue set based on gene-centered networks obtained for all the genes present in the grape gene reference catalogue. TI versus berry (top panel), TI versus leaf (middle panel) and berry versus leaf (bottom panel). Scatterplots were generated using only co-expression relationships shared between the compared tissue sets.

### GCNs as TF-function prediction tools

Single gene-centered networks of each member of the R2R3-MYB and bHLH gene families (134 and 95 genes respectively) were generated. Each gene-centered network contained only the top 420 most co-expressed genes with the gene of interest. Individual gene-centered networks were then subjected to a gene set enrichment analysis (GSEA) based on hypergeometric distribution and false discovery rate (FDR) pvalue adjustment, performed using gprofiler2 R package (Kolberg et al., 2020) using the manually curated MapMan ontology (Sup. Dataset 2). R2R3-MYBs and bHLHs presenting significant enrichment (FDR < 0.05) of at least one MapMan bin related with the secondary metabolism were selected as potential regulators.

### Validation of GCNs using cistrome data

GCNs were overlapped with TF genome-wide binding data to evaluate their capacity to predict regulatory networks. MYBPA1 DNA Affinity Purification Sequencing (DAP-Seq) was performed as detailed in (Orduña et al., 2022), DNA-binding motif was generated by running the MEME suite 5.5.0 (Machanick & Bailey, 2011) with default parameters on the sequence of the top 600 most significant binding events (i.e., peaks). Binding motifs comparison were executed with TFBSTools R package (Tan & Lenhard, 2016). DAP-Seq bound genes were overlapped with the gene-centered networks derived from berry, leaf and TI filtered aggregated GCNs and common genes were defined as berry, leaf and TI high-confidence targets (HCTs). HCTs were validated *in planta* by overlapping them with a list of differentially expressed genes in response to MYBPA1 overexpression in hairy roots, in order to generate berry, leaf and TI very high confidence targets (VHCTs) lists.

### Microarray probe re-annotation

The Qiagen Operon Array-Ready Oligo Set for the Grape Genome Version 1.0, used in (Terrier et al., 2009), contained 14,562 70-mer probes representing 14,562 transcripts. Data from the microarray analysis was downloaded, and probe-to-gene associations were reassigned by performing a blastn of the probe sequences with VCost.v3_27 transcripts using default parameters. In case of probes assigned to more than one gene, assignments were filtered according to their blastn evalue, keeping only the association with the lowest evalue. In the case of a probe assigned to more than one gene with the same evalue, both associations were considered valid. Probes with upregulation in (Terrier et al., 2009) were extracted and converted to VCost.v3 gene IDs using the previously generated associations.

## RESULTS

### Network analysis

Transcriptional activators represent an excellent case study to test and validate the efficacy of gene co-expression networks (GCNs) as their targets tend to closely follow activator expression patterns. These relationships, however, may be dependent on certain conditions, e.g., in response to specific stimuli or in precise organs, tissues or developmental stages. Thus, we inspected gene co-expression relationships among all genes present in the grapevine genome by constructing tissue-independent (TI) and dependent GCNs.

To generate the GCNs, 4,815 SRA runs (i.e., libraries) were downloaded from the Sequence Read Archive (SRA) database. After the manual and automatic filtering process, GCNs were built from 67 SRA studies (1,615 SRA runs), 42 SRA studies (671 SRA runs) and 131 SRA studies (2,767 SRA runs) for berry, leaf and tissue-independent (TI) GCNs, respectively (Table 1). Two different aggregation approaches were used to build each network, leading to aggregated (i.e., co-occurrence-based aggregation) and single (a network constructed from merged datasets) GCNs. In essence, our GCNs report 1% of the most co-expressed genes in the genome for each gene (a list of 420 genes, since the PN40024 12X.v2 assembly has 42,413 genes), ordered by interaction value (i.e., the co-occurrence). The number of nodes and edges contained in each generated network is always comparable between the aggregated and single approaches (Table 2).

**Table 1.** General overview of transcriptomic datasets used for GCNs construction. Automatic and manual SRA run filtering reduces the number of transcriptomic datasets available for the GCNs generation.

| **Tissue** | **Unfiltered SRA runs** | **SRA Runs used in GCNs construction** | **SRA studies used in GCN construction** |
| --- | --- | --- | --- |
| leaf | 1210 | 672 | 42 |
| fruit | 2,223 | 1615 | 67 |
| root | 61 | 36 | - |
| stem/shoot | 62 | 32 | - |
| woody stem | 83 | 69 | - |
| flower | 184 | 159 | - |
| seed | 60 | 55 | - |
| tendrill | 3 | 3 | - |
| bud | 207 | 96 | - |
| Tissue mix | 71 | 29 | - |
| **TOTAL (TI)** | **4164** | **2766** | **131** |
| unclassifiable | 651 | 648 |  |
| **TOTAL** | **4815** | **3414** |  |

**Table 2.** Basic GCNs formal analysis. Network parameters are significantly affected by a combination of tissue set choice and aggregation approach.

| Network | Number of nodes | Number of edges | MapMan AUROC value | GO AUROC value |
| --- | --- | --- | --- | --- |
| **Berry aggregated GCN** | 30,831 | 12,949,020 | 0.7527 | 0.6892 |
| **Berry single GCN** | 30,919 | 12,985,980 | 0.7642 | 0.7129 |
| **Leaf aggregated GCN** | 31,759 | 13,338,780 | 0.7689 | 0.7010 |
| **Leaf single GCN** | 32,174 | 13,513,080 | 0.7669 | 0.7057 |
| **TI aggregated GCN** | 35,732 | 15,007,440 | 0.7924 | 0.7228 |
| **TI single GCN** | 36,082 | 15,154,440 | 0.7877 | 0.7265 |

A GCN can be envisaged as a classification system based on the *guilt-by-association* principle, whereby co-expressed genes are expected to share similar biological roles (Wolfe et al., 2005). Because of this, GCN performance is typically assessed through gene function prediction metrics based on gene ontologies such as the area under the receiver operating characteristic curve (AUROC) parameter: an indicator of the probability that a gene will cluster with the rest of the genes of its ontology term. Thus, the performance of our networks was measured using the AUROC parameter by means of a threefold cross-validation across manually curated MapMan V4 BIN functional categories and Gene Ontology (GO) terms associated to the VCost.v3 annotation. To ensure robustness, only MapMan bins and GO terms with 20-1000 annotated genes were used for AUROC performance evaluation. AUROC values were computed across different numbers of studies cumulatively (Sup. Figure 2) to study the impact of dataset size in both types of pipelines. The observed AUROC values were more sensitive to the selection of experiments with subset sizes below 30 regardless of tissue set (leaf, berry or the tissue-independent dataset), network building approach (aggregated or single), and ontology type (MapMan or GO). Conversely, above 30 experiments, AUROC values tend to stabilize and additional experiments result in smaller improvements in network performance (Figure 1A).

Once all SRA studies were added to their respective GCNs, clear differences when comparing AUROC values between tissues, ontologies used for evaluation and different methodologies were shown. In the first place, the MapMan ontology always yielded a higher AUROC compared to GO for the same GCN. Secondly, when using MapMan, the aggregated approach yielded higher AUROC values with respect to the single approach in the TI and leaf GCNs, whereas the single approach outperformed the aggregated approach in the berry GCN.

To test the impact of the two aggregation methods in the AUROC of the GCNs, a network was generated from each individual SRA study, and their AUROCs were computed using MapMan and GO ontologies. The highest AUROC values for individual SRA study networks using MapMan were 0.7272, 0.7488 and 0.7508, and 0.6772, 0.6951 and 0.6994 using GO for berry, leaf and TI experiments respectively, whereas the lowest values were 0.5677, 0.5669 and 0.5678 for MapMan and 0.5414, 0.5395, and 0.5458 for GO (Sup. Table 2). These values show that independently of the tissue analyzed, both aggregation approaches (aggregated and single) improve network performance when compared to individual SRA studies (Figure 1B & Sup. Figure 3).

**Figure 3.**
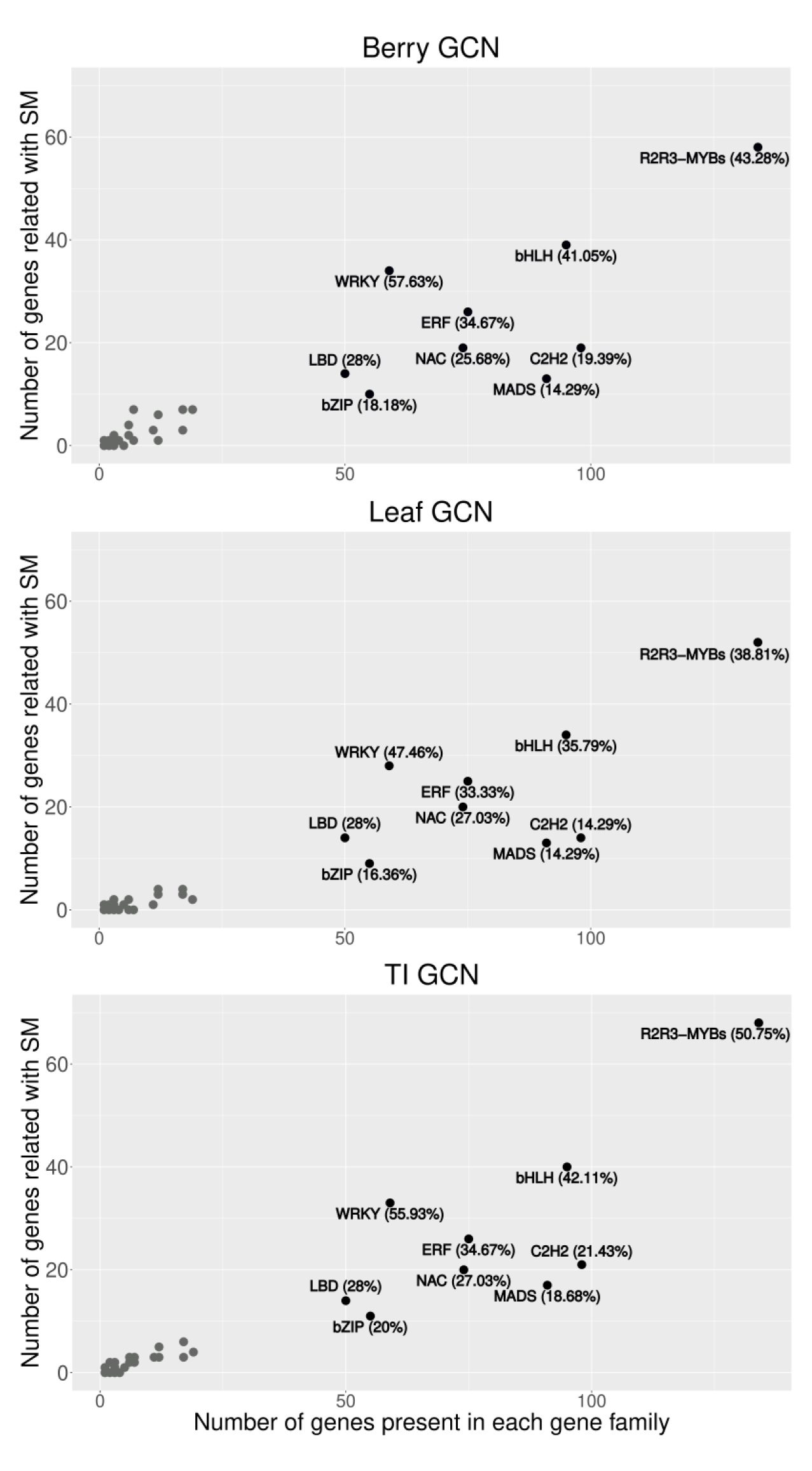
Screening of the potential involvement in the secondary metabolism regulation for all TF families found in the grape gene reference catalogue. Gene families with more than 50 members appear highlighted in black color, while grey dots represent the following TF gene families: ARF, SBP, TCP, AP, GRAS, YABBY, RAV, PRR, HCBT, ELF, RGA, WDR, TOC, HD, ZIP, Gigantea, LHY, LUX, RVE and PIF.

### GCNs as tissue-specific gene function prediction tools

To further analyze the differences between GCNs coming from different tissue sets and their use as TF-function prediction tools, we generated gene-centered networks across each gene present in the grape gene reference catalogue, a repository of all available information on characterized genes in grapevine (Navarro-Payá et al., 2022). Gene-centered networks were derived from the aggregated GCNs from the three analyzed tissue sets. These gene-centered networks contained the 420 most co-expressed genes with the gene of interest. The co-occurrence values of shared co-expression relationships between the chosen tissue sets for network construction was compared in two-dimensional scatterplots (Figure 2). Co-expression values were normalized according to the number of SRA studies each GCN was built from (131, 67 and 42 for TI, berry and leaf respectively). The linear regression coefficient (R^2^) values indicate a high similarity between the TI-berry and TI-leaf comparisons. Nevertheless, the berry-leaf comparison has a lower R^2^ value (0.825) suggesting these are the more divergent tissue sets in terms of co-expression relationships. This fact implies that tissue-specific GCNs performance may be significantly different across the different MapMan bins, since tissue-specific GCNs are able to detect tissue-specific co-expressions that may be biologically relevant. The value of AUROC of each MapMan bin cannot be compared between the three GCNs because the number of datasets from which they are built is not the same. However, if it is scaled to z-score, it can be used to compare all GCNs. For instance, MapMan bin 27.1, related to the circadian clock system, has higher z-score AUROC value in the leaf GCN (1.3338) when compared to the berry GCN (−0.625). On the contrary, MapMan bin 5.9, related to lipid metabolism, has a higher z-score AUROC in the berry GCN (0.8243) than in the leaf GCN (−1.3542). Regarding tissue-independent GCN, AUROC values across these MapMan bins are 0.2095 and 0.4427, respectively (Sup. Table 1 & Sup. Figure 4).

**Figure 4.**
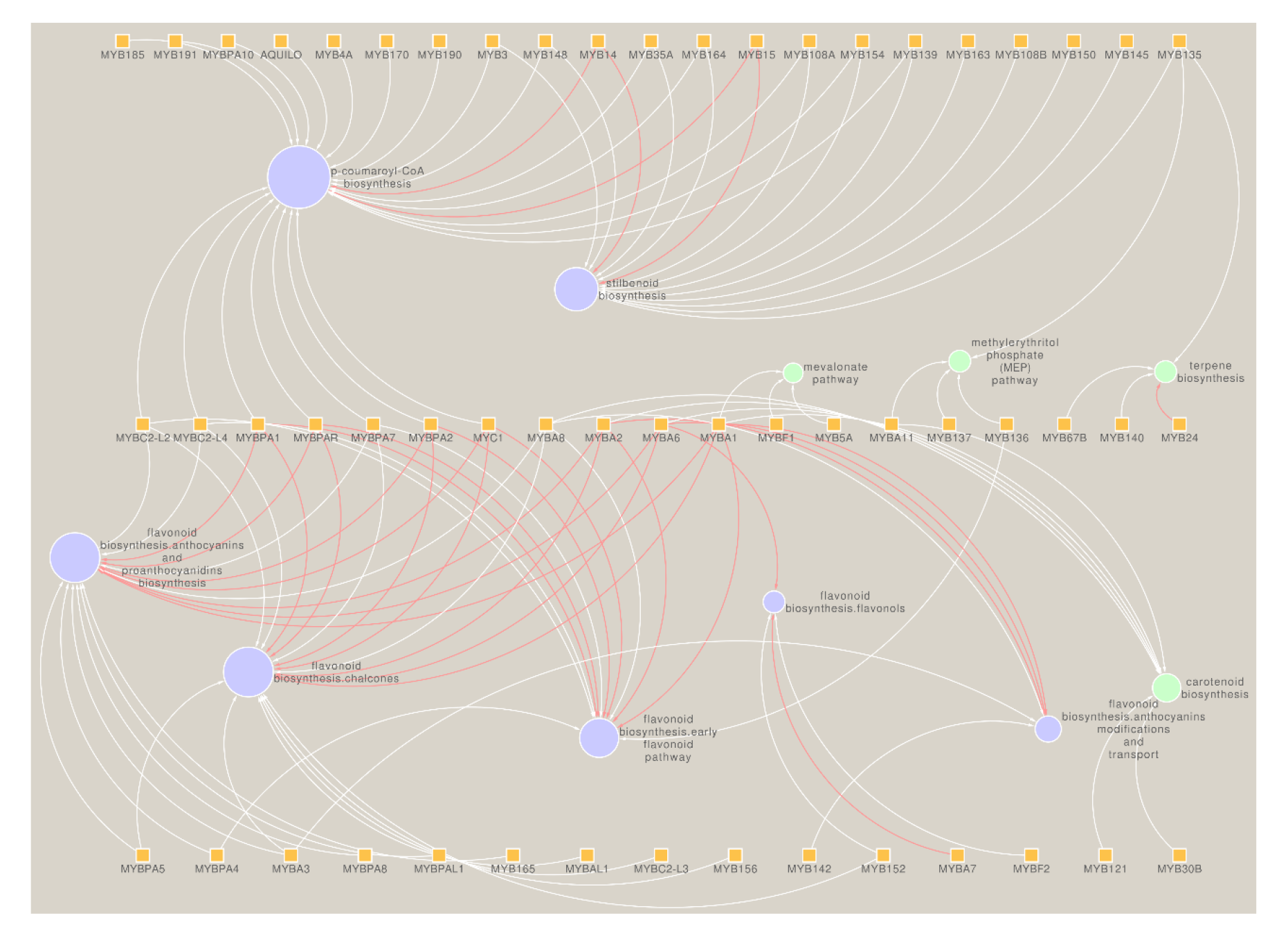
MYB gene family known regulatory roles are reflected in the leaf secondary metabolism network, while potential new regulatory roles are suggested. TI MYB family secondary metabolism network, that includes all R2R3- MYB genes (square nodes) that present at least one secondary metabolism related bin significantly enriched on its gene-centered network enrichment analysis. Circles depict MapMan bins involved in the secondary metabolism, and their size are delimited by the number of MYB genes whose gene-centered networks are enriched in the respective MapMan bin. Red edges depict regulatory relationships already described by literature.

Additionally, for each gene present in the grape reference catalogue, a 3D scatterplot was generated representing co-occurrence values across the TI, berry and leaf filtered aggregated gene-centered networks, respectively. All generated 3D plots are available at the 3Dplots app at www.vitviz.tomsbiolab.com.

After testing differences in co-expression between the analyzed tissue sets, we explored whether these RNA-seq derived gene co-expression networks could serve as a means to propose roles of uncharacterized genes. If GCNs serve as predictors, then we were also interested in understanding which fraction of a network holds a close and true functional relationship to a gene of interest. As a proof of concept, and considering TF bins such as WRKY, bHLH, MYB-related and bZIP bins generally perform better for AUROC calculation in aggregated networks (Sup. Table 2), we evaluated gene-centered networks extracted from the aggregated GCNs (for the three tissue-sets) of all so-far described transcriptional regulators found in the grape gene reference catalogue (version 2, (Navarro-Payá et al., 2022). It is possible to determine, by using different experimental validation methods, if genes that belong to a GCN are actual targets of a given TF.

First, an enrichment analysis using manually curated MapMan ontology was done across each one of the previously generated gene-centered networks, keeping only genes whose networks present at least one statistically significant enrichment for a function related to secondary metabolism. All TF gene families included in the grape gene reference catalogue were screened, identifying the total number of family members that seem related to secondary metabolism. Across the berry, leaf and TI GCNs, the R2R3-MYB gene family presented the highest number of members potentially related to different pathways and branches of secondary metabolism, followed by the bHLH gene family. The WRKY gene family, interestingly, presented the highest percentage of members potentially involved in secondary metabolism across the three evaluated GCNs (Figure 3).

Since the R2R3-MYB and bHLH gene families were the ones with more members potentially involved in the secondary metabolism according to the three tissue sets GCNs, further analysis was carried out on them. For the berry GCN, out of the 208 R2R3- MYBs and bHLH genes considered (120 and 88 genes with expression data, respectively), 65 (41 MYBs and 24 bHLHs) were enriched in secondary metabolism categories, while in leaf GCNs (209 MYBs/bHLH analyzed) 69 (42 MYBs and 27 bHLHs) were enriched in these terms. In the TI GCN, the ratios slightly increased, with 80 TFs enriched in the terms of interest (55 MYBs and 25 bHLHs), out of 226 being analyzed.

Our methodology was able to recover previously described roles of characterized TFs, e.g., MYB14/15 being involved in the regulation of p-coumaroyl-CoA and stilbenoid biosynthesis (Orduña et al., 2022), MYBA1/A2/A6 involved in all the steps of the anthocyanins biosynthesis (Matus et al., 2017; Rinaldo et al., 2015; Walker et al., 2007), and *MYC1* involved in different steps of proanthocyanidin biosynthesis (Hichri et al., 2010). These relationships were conserved across berry, leaf and TI GCNs. Additionally, new potential regulatory roles that should be experimentally confirmed, were also suggested (i.e., MYBPA1 putatively involved in *p*-coumaroyl-CoA biosynthesis, or MYB108A related to the stilbenoid biosynthesis, among other phenylpropanoid branches. Furthermore, some significant relationships between MYB/bHLH genes and secondary metabolism categories (bins), such as the ones previously mentioned, were conserved across the berry, leaf and TI networks, while others resulted tissue-specific (i.e., MYB108B is inferred as related to *p*-coumaroyl-CoA biosynthesis in the leaf GCN and with anthocyanin transport and modification in the berry GCN). This suggests that the different GCNs are able to detect organ-specific relationships according to the samples used in their generation (Figure 4, Sup. Figure 5, Sup. Figure 6, Sup. Table 4).

**Figure 5.**
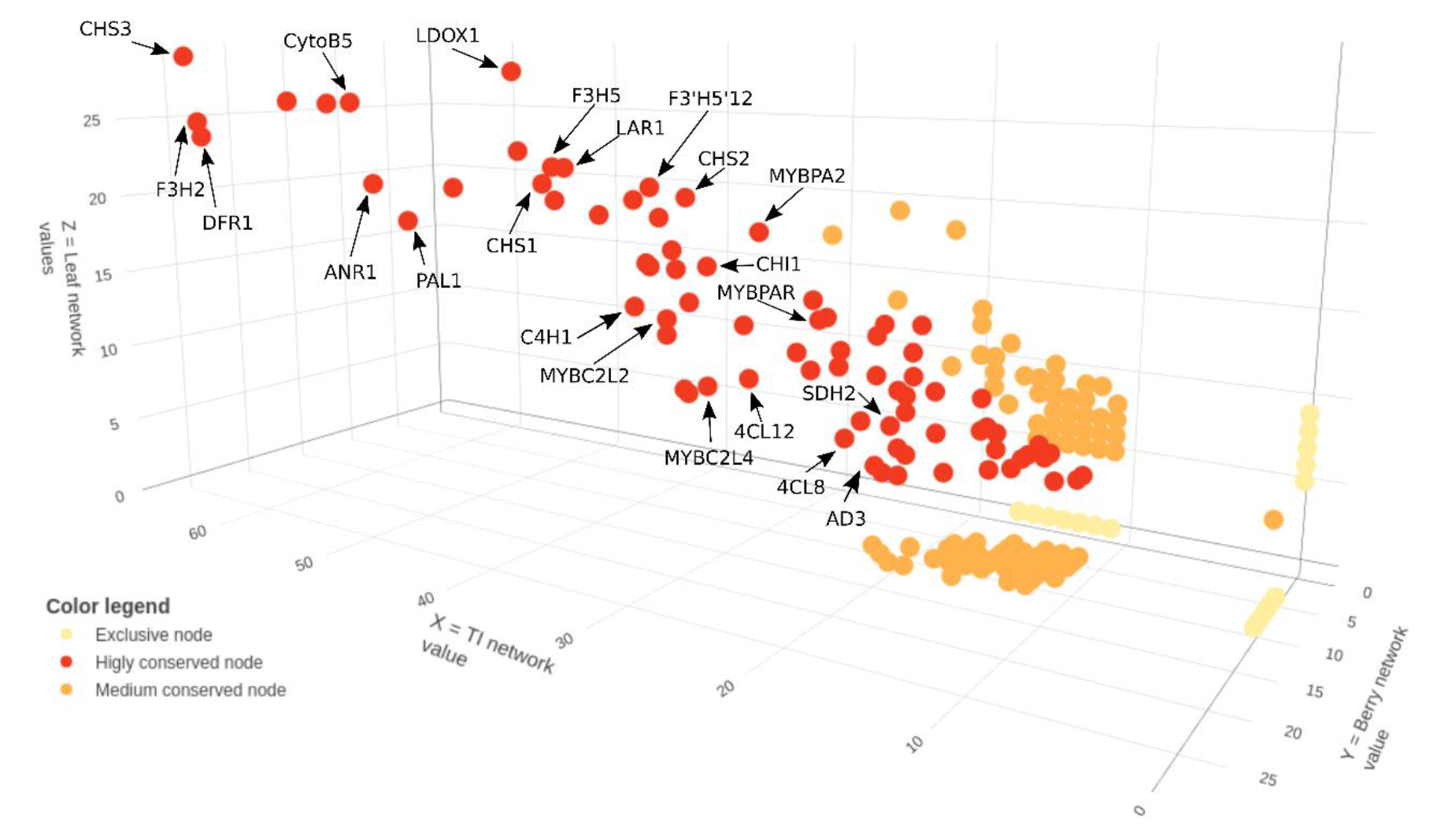
Shared co-expressed genes between TI, berry and leaf gene-centered networks show MYBPA1 as highly connected with phenylpropanoid and proanthocyanidin metabolism. MYBPA1 common nodes across the three tissue sets share high co-occurrence values and mostly represent different steps and branches of the phenylpropanoid and proanthocyanidin pathways (marked by black arrows), including both structural genes (PAL1/ANR1/CHS1) and their regulators (MYBPA2/MYBC2L2). MYBPA1 3D plot, where the X, Y and Z axis depict the MYBPA1 gene-centered network co-occurrence values for the TI, berry and leaf GCNs, respectively.

**Figure 6.**
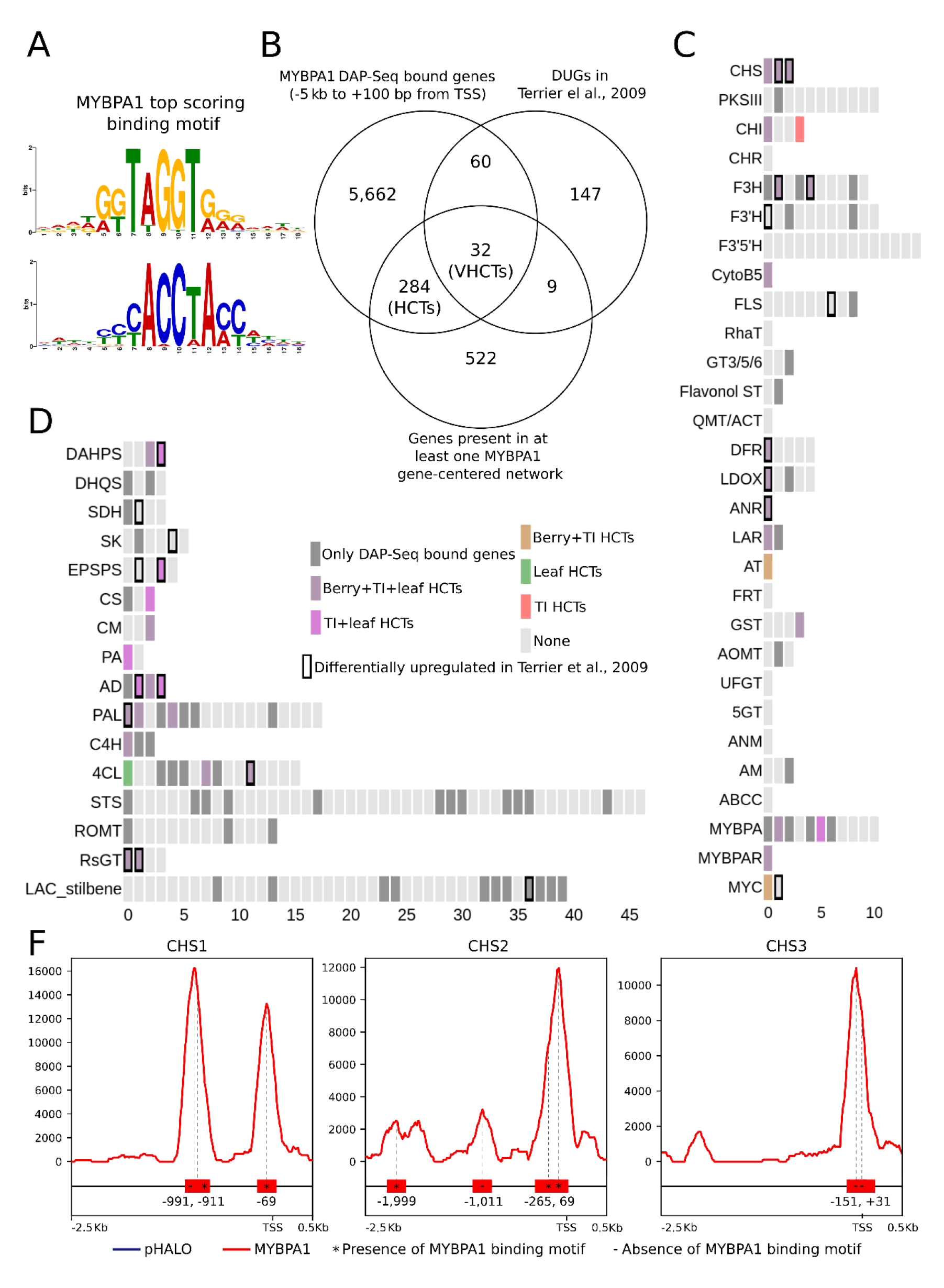
Identification of new MYBPA1 high confidence targets based on the overlap between MYBPA1 DAP-Seq, MYBPA1 gene-centered networks and MYBPA1 overexpression transcriptomic dataset. A) *De novo* binding motifs, forward and reverse, obtained from the top 600 scoring peaks of MYBPA1 DAP-Seq using MEME suite with default parameters. B) Venn diagram of shared genes between DAP-Seq bound genes within −5 kbp to +100 bp to the TSS, the genes present in the TI, leaf and berry gene-centered networks and the DUGs in the microarray analysis performed in (Terrier et al., 2009). C) Genes from the flavonoid and D) shikimate, early phenylpropanoid and stilbenoid pathways are sorted from left to right and top to bottom, with different color legend according to its classification as MYBPA1 DAP-Seq bound genes or HCTs according to the different tissues GCN. VHCTs are highlighted with black borders. Full gene names of the genes shown in these pathways are available at (Sup. Dataset 3). F) MYBPA1 DAP-Seq binding signal in the promoter regions of the chalcone synthase (CHS) gene family members. Binding events that have MYBPA1 binding motif are shown with an asterisk.

### *MYBPA1* as a regulator of flavonoid and *p*-coumaroyl-CoA biosynthesis

According to MapMan enrichment analysis of the berry, leaf and TI gene-centered networks, *MYBPA1* is related to the regulation of chalcone synthase activity and proanthocyanidin (PA) biosynthesis, two roles previously demonstrated for this TF (Bogs et al., 2007; Terrier et al., 2009). Additionally, in the enrichment analysis, MYBPA1 is also closely related in expression to the early flavonoid pathway, the anthocyanin branch and also *p*-coumaroyl-CoA biosynthesis the latter occurring at the initial steps of the phenylpropanoid pathway. A closer inspection of the MYBPA1 gene-centered networks in the form of a 3D scatterplot reveals that important genes involved in shikimate (*DAHPS3*, *SK5*), *p*-coumaroyl-CoA (*PAL1/2/5*, *C4H1* and *4CL8/12*) and proanthocyanidin (*CHS1/2/3*, *CHI1*, *F3H2/5*, *F3’H1/2*, *CytoB5*, *DFR1*, *LDOX1*, *ANR1*, *LAR1* and *3AT*) biosynthesis are shared with high co-occurrence values between all tissue conditions (Figure 5). In addition, interesting TF-coding genes also appear in the three MYBPA1 gene-centered networks, such as MYBPA2 (Terrier et al., 2009) and MYBPAR (Koyama et al., 2014), and the C2 clade repressors *MYBC2L2* or *MYBC2L4* (Cavallini et al., 2015).

To further corroborate the previously asserted roles of MYBPA1 in PA regulation and demonstrate the newly proposed ones, DNA affinity purification sequencing (DAP-Seq) was carried out, with MYBPA1 being amplified from cv. ‘Pinot Noir’ (PN), challenged with cv. PN gDNA, and reads being mapped to the PN40024 12X.2 assembly. Our DAP-Seq reported 8,109 MYBPA1 binding events (i.e., peaks) present between 5 kb upstream to 100 bp downstream of the transcription start site (TSS, out of 30,728 total binding events), which were associated with 6,038 VCost.v3 gene models in total (Sup. Table 3). The top 600 most significant peak sequences, used for obtaining the DNA-binding motif, show that the MYBPA1 mainly binds to the “GGTAGGT” consensus sequence (Figure 6 A). The detected binding motif is very similar to the binding motifs found for MYB14/15 (Orduña et al., 2022), with Pearson correlation coefficients of 0.916 and 0.91 for MYB14 and MYB15 respectively (Sup. Figure 7).

**Figure 7.**
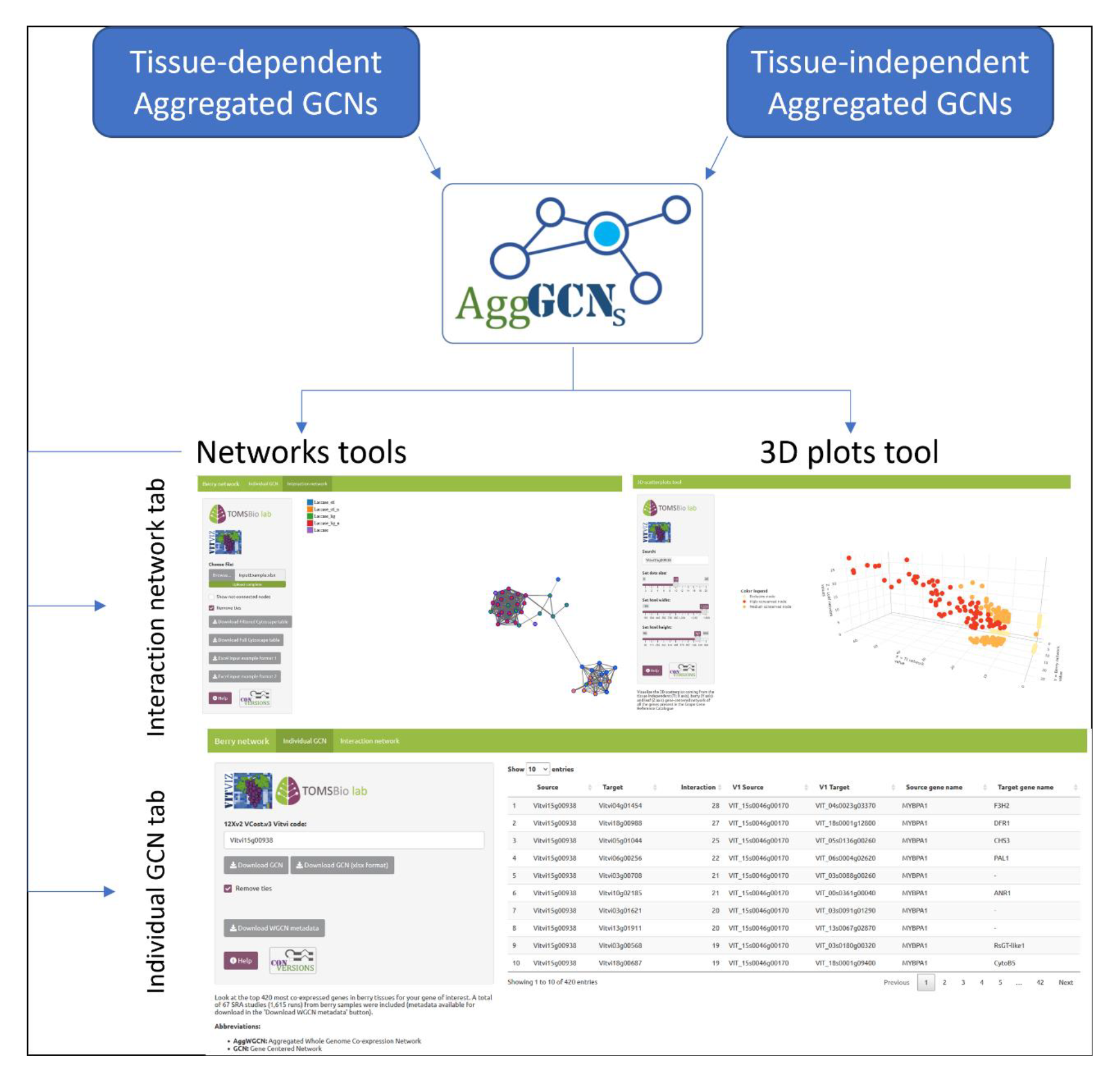
AggGCNs app allows user-friendly exploration of the GCNs and 3D plots. The AggGCNs app takes as input both the tissue-dependent and independent GCNs. From that, it allows the user to either access the 3D plots tool, where it is possible to generate the 3D plot for each gene present in the grape gene reference catalogue. On the other hand, users can also select a particular GCN network tool, where they can generate the gene-centered network of a gene or their interest in the individual GCN tab. A list of genes of interest can also be uploaded to generate an interactive network with all the connections present within the list of genes that the user has provided across the GCN in the interaction network tab.

A closer look at the DAP-Seq peaks revealed that a high number of genes involved in the shikimate and early phenylpropanoid pathways that ultimately lead to biosynthesis of p-coumaroyl-CoA, are bound by MYBPA1, supporting our previous observations extracted from the MYBPA1 gene-centered networks. Out of the 35 genes involved in the shikimate and aromatic amino acid pathway, 15 are MYBPA1 DAP-Seq bound genes (the 3-deoxy-D-arabino-heptulosonate 7-phosphate synthases *DAHPS3-4*, the 3- dehydroquinate synthases *DHQS1* and *DHQS3*, the shikimate dehydrogenase *SDH1*, the shikimate synthase *SK2*, the 5-enolpyruvylshikimate 3-phosphate synthase *EPSPS4*, the chorismate synthases *CS1* and *CS3*, the chorismate mutase *CM3*, the prephenate aminotransferase *PA1*, and the arogenate dehydratases *AD1-4*), while in the case of the early phenylpropanoid pathway, out of the 37 genes involved, 17 were bound by MYBPA1 (the phenylalanine amonio lyases *PAL1*-*2*, *PAL4*-*7* and *PAL14*, the Cinnamate-4-hydroxylases *C4H-3*, and the 4-coumarate:CoA ligases *4CL1*, *4CL4*-*6*, *4CL8*-*9* and *4CL12*). On the other hand, MYBPA1 DAP-Seq results also corroborate the relationship between MYBPA1 and the biosynthesis of proanthocyanidins, since 19 key genes involved in this pathway were bound by MYBPA1 (the chalcone synthases *CHS1*-*3*, the chalcone-flavonone isomerases *CHI1* and *CHI4*, the flavanone 3-hydroxylases *F3H1*-*2*, *F3H4*-*5* and *F3H9*, the flavonoid 3’ hydroxylases *F3’H3* and *F3’H9*, the cytochrome p450 *CytoB5*, the dihydroflavonol 4-reductase *DFR1*, the leucoanthocyanidin dioxygenases *LDOX1* and *LDOX3*, the anthocyanidin reductase *ANR1*, and the leucoanthocyanidin reductases *LAR1-2*). Interestingly, MYBPA1 binds itself at the upstream regulatory region.

MYBPA1 bound genes were overlapped with the MYBPA1 gene-centered networks extracted from the berry, leaf and TI filtered aggregated GCNs (Sup. Table 3), generating a dataset of high confidence targets (HCTs) present in at least one tissue MYBPA1 gene-centered network. A total number of 316 genes were described as HCTs (Sup. Table 3). Nine (*DAHPS3-4*, *ESPSP4*, *CS3*, *CM3*, *PA1*, *AD2-4*), 7 (*PAL1-2*, *PAL5*, *C4H1*, *4CL1*, *4CL8* and *4CL12*) and 13 (*CHS1-3*, *CHI1*, *CHI4*, *F3H2*, *F3H5*, *CytoB5*, *DFR1*, *LDOX1*, *ANR1*, *LAR1* and *3AT1*) HCTs are present in the shikimate, early phenylpropanoid and proanthocyanidin biosynthesis pathways, respectively. In addition, *MYBPA2*, *MYBPAR* and *MYC1*, TFs that are known as PA-positive regulators, were also detected within the HCT list.

In order to validate the HCTs *in planta*, we reanalyzed the differentially upregulated genes detected in (Terrier et al., 2009), where authors overexpressed *MYBPA1* in *V.vinifera* hairy roots. The authors analyzed the transcriptome of the *MYBPA1* overexpressing hairy roots with the Qiagen Operon Array-Ready Oligo Set for the Grape Genome Version 1.0, which contained 14,562 70-mer probes representing 14,562 transcripts. Our microarray probes re-annotation pipeline was able to recover 248differentially upregulated genes (DUGs) VCost.v3 gene IDs, that were overlapped with the previously detected HCTs to obtain the final ‘very high confidence targets’ (VHCTs). 32 VHCTs were detected by this overlap (Figure 6 B, Sup. Table 3). From these, three VHCTs are involved in the shikimate pathway (*DAHPS4*, *EPSPS4* and *AD2*), while two VHCTs are involved in the early phenylpropanoid pathway (*PAL1*, and *4CL12*) and seven more VHCTs are involved in proanthocyanidin biosynthesis (*CHS2-3*, *F3H2*, *F3H5*, *DFR*, *LDOX1* and *ANR1*; Figure 6 C & D).

Based on these results, we thus confirm that the GCNs were able to predict already known MYBPA1 regulatory interactions with key genes of the PA biosynthesis pathway, such as *ANR1*, *LAR1-2* and *LDOX1*. In addition, GCNs were also able to predict MYBPA1 regulatory interactions with more genes of the flavonoid and early phenylpropanoid pathways, which were confirmed by overlapping MYBPA1 gene-centered networks with DAP-Seq and microarray datasets. For instance, chalcone synthase (*CHS*) genes, that catalyze the first step of the flavonoid pathway, preceding the biosynthesis of proanthocyanidins, and as *CHS1* and *CHS2/3,* appeared as HCT and VHCTs, respectively, with clear MYBPA1 binding in their promoter sequence (Figure 6 F).

### The aggGCN application: an online user-friendly app for navigating through GCNs

To access and navigate through the information contained in the generated GCNs, the aggGCN app was developed within the context of the VitViz platform (http://www.vitviz.tomsbiolab.com/). This app introduces several user-friendly functionalities for analyzing both the tissue-dependent and tissue-independent GCNs (Figure 7). Upon accessing the app, users can choose between the Gene Centered 3D Plots tool or a particular GCN. If the Gene Centered 3D Plots tool is selected, users are able to introduce the VCost.v3 gene ID of their interest, and the tool will automatically generate an interactive 3D scatterplot of the gene of interest.

On the other hand, if the user selects the tool of a particular GCN (i.e., TI GCN), a new window with two tabs is displayed. The “Individual GCN” tab allows users to generate and download gene-centered networks from any gene of interest from the aggregated GCN. Since it is possible to have ties among the lowest co-occurrence values on each network while selecting the top 420 most highly co-expressed genes, two versions of the gene-centered network of the gene of interest were generated. On a first version, ties were not removed, and therefore gene-centered networks contain variable gene size, while in the second version, ties have been removed according to their order of appearance (based in ID) in the GCN matrix. These later networks always contain 420 genes, and are the ones used for the analysis described in this study.

The “Interaction network” tab allows users to upload a list of genes of interest and generate an interactive network with all the co-expression relationships present in that gene set, across the aggregated GCNs, also giving the user the choice of keeping or removing ties in the lowest co-expression values. These interaction plots represent a suitable way to visualize a pathway-oriented network, where the user can inspect how genes present in the gene set given as input are clustered according to their co-expression values. For instance, this tool can be used for separating the *STS* gene family and their known regulators (*MYB14-15*) from the proanthocyanidin (*ANR1* and *LAR1-2*) and anthocyanin biosynthesis (*UFGT1*, *AOMT1*, *GST4* and *3AT*) key-pathway genes and their respective known regulators, adding uncharacterized TFs in order to predict the biosynthetic pathway they may be regulating (Figure 8).

**Figure 8.**
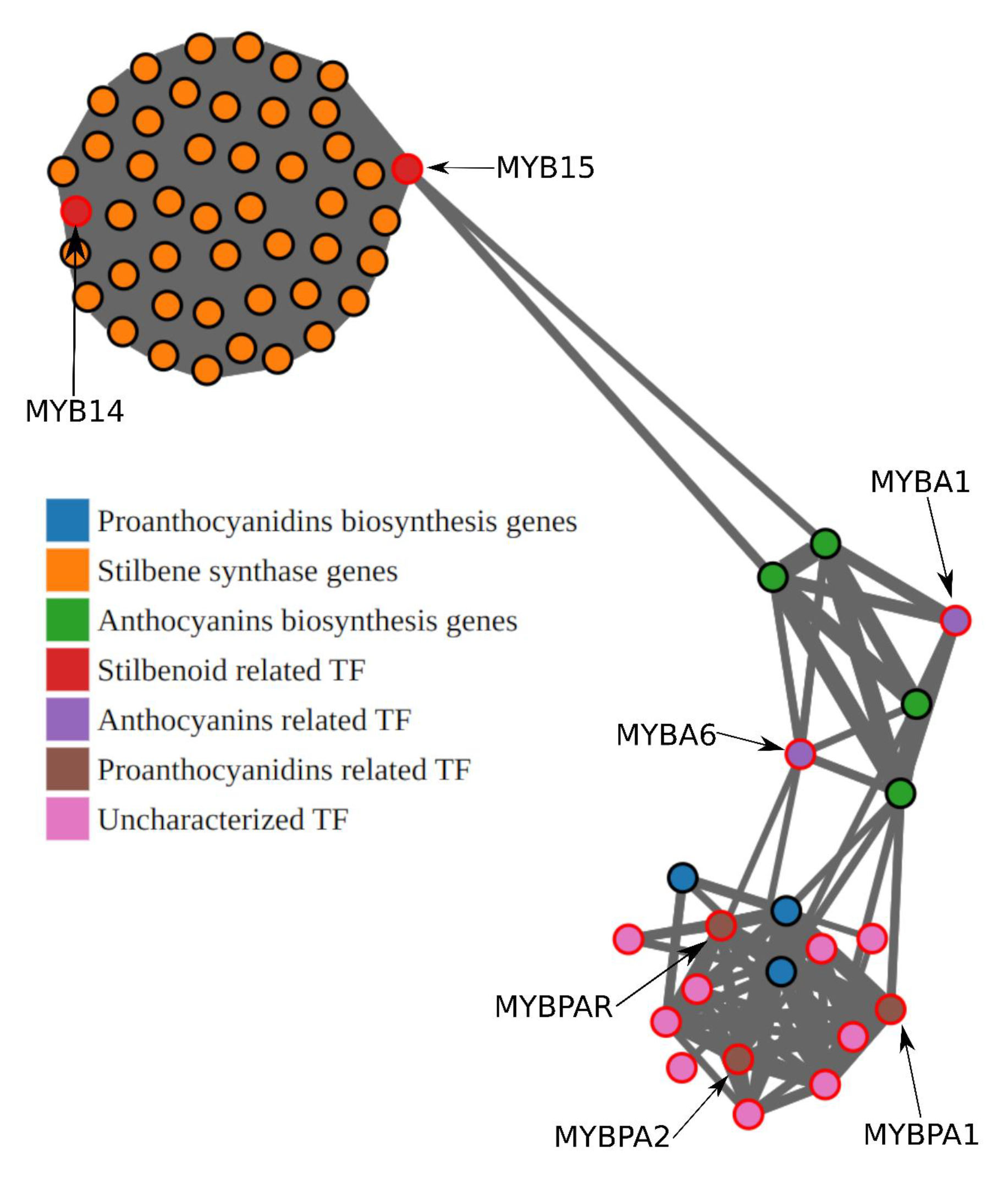
Interaction plots are able to cluster genes according to the co-expression, providing a new way of identifying potential pathway regulators. TFs (nodes with border highlighted in red) cluster with the genes they are regulating. Uncharacterized TFs (*MYC2-L2*, *MYBC2*-*L4*, *MYBPA4-5*, *MYBPA8*, *MYBPAL1*, *MYB165*, *MYB156* and *MYB152*), that appear as related with the proanthocyanidin pathway in Figure 4, also appear related with the proanthocyanidin biosynthesis genes (*ANR1* and *LAR1-2*) and known proanthocyanidin regulators (*MYBPA1-2* and *MYBPAR*), whereas *MYBA1* and *MYBA6*, known anthocyanins regulators, appear related with anthocyanins biosynthesis genes (*UFGT1*, *AOMT1*, *GST4* and *3AT*).

## DISCUSSION

### Aggregation improves network performance

The present study comprises a deep analysis of the performance of different network construction methods, involving aggregation and condition dependency, some of which have recently been demonstrated to be useful in the characterization of grapevine TFs such as *NAC60*, *MYB14*, *MYB15* and *MYB24* and (D’Incà et al., 2023; Orduña et al., 2022; Zhang et al., 2021). Network performance was assessed with the information contained in the grapevine MapMan and GO ontologies using the area under the receiver operating characteristic curve (AUROC), which is a measure of the degree to which network connectivity can predict membership within a given gene set defined by functional ontology terms (Ballouz et al., 2017).

In agreement with previous studies (Ballouz et al., 2015; Verleyen et al., 2015), our evaluation of network performance (at least in the ways tested in this work, i.e., co-occurrence aggregation or single network construction from merged datasets) highlighted that aggregation performs better than networks constructed from individual SRA studies as no single SRA study network outperformed any of the generated GCNs independently of the ontology used. Nevertheless, remarkable differences appear in the AUROC parameter when comparing the ontology (i.e., MapMan or Gene Ontology GO) used for the assessment and the performance between aggregation approaches. In the first place, AUROC values can be biased if relationships contained in the used ontology do not reflect real biological relationships, retrieving different AUROC values for the same GCN. For example, the stilbenoid biosynthesis pathway bin was manually curated because several stilbene synthases (STS), stilbenoid-related laccases and resveratrol-O-methyl transferases had not been previously included (the same occurred for anthocyanin-modifying enzymes). Here, we observe that MapMan systematically yields a better AUROC value than GO for the same networks (Figure 1A & Table 2). This can be explained because MapMan a1nnotation was designed for plant-specific processes, containing less ambiguous terms (Thimm et al., 2004). Thus, MapMan AUROC measurements seem more reliable than GO measurements, corroborating previous studies (Klie & Nikoloski, 2012).

While comparing aggregation approaches, co-occurrence-based aggregation generally outperforms single network construction from merged datasets. This occurs when using all available SRA studies for each GCN, together with the assessment of the MapMan ontology. The only exception was the berry GCN, where the single approach has a higher AUROC value. On the other hand, when using GO, the single approach slightly outperformed the aggregated method. These AUROC differences are consistent across all MapMan bins and GO terms (Figure 1B & Sup. Figure 3), meaning that network performance measurements are not biased by any particular category.

Altogether, and because MapMan is a more consistent ontology, our results suggest that aggregated approaches generate more reliable GCNs when using an initial dataset of at least 20 SRA studies. This has also been suggested when using microarray datasets (Huang et al., 2017; Liesecke et al., 2019; D. C. J. Wong, 2020). The AUROC value from the single approach is slightly higher in the berry GCN, and this could be explained by the fact that berry transcriptomes contain specific co-expressions (e.g., tissue-, stress-or cultivar-related) that influence the overall variability of berry relationships (Dal Santo et al., 2013). Although MapMan is the best ontology adapted to grapevine, it may not be able to correctly capture these additional specific co-expressions because some bins are incomplete or contain an incorrect set of genes. A good example explaining this is given by the differences between red-skin and white-skin cultivars. In red-skins cultivars, the genes of the *chalcone synthase* (*CHS*) family and the *flavonoid 3-O-glucosyltransferase* (*UFGT1*) are co-expressed, because they are involved in anthocyanin synthesis. However, in white-skin cultivars they are not co-expressed since *UFGT1* is not expressed (its regulator MYBA1 is knocked-out by the GRET1 transposon insertion), while *CHS* genes are being transcribed. In the MapMan ontology, the 9.2.2 bin, which contains the genes involved in the flavonoid metabolism, both *CHS* genes and the *UFGT1* are included. Therefore, depending on the cultivar, the MapMan ontology bin will be more or less accurate. For example, through the comparison of white and red cultivars, (Massonnet et al., 2017) was able to identify a core-set of approximately 7,000 genes potentially regulated (cause and/or effect) by the presence/absence of anthocyanins in berry skins. Aggregation may introduce noise when merging the red and white cultivar datasets, with the noise likely coming from these 7000 genes.

We clearly observe that network performance is affected by the number of studies used. When using a reduced number of SRA studies, performance of aggregated approach results is highly variable between subset sizes and iterations (as seen in the size of the boxplots in Figure 1A). We suggest that a robust aggregated network can be constructed with at least 25-30 SRA studies. This would be the first requirement before implementing a GCN approach on a non-model species.

Absolute AUROC or co-occurrence values are not comparable between GCNs generated from different tissue sets. This occurs because GCNs are constructed using datasets of different sizes (Table 1). This issue is overcome when representing co-occurrence values in percentages or by scaling (e.g., z-scored) AUROC values as we show that, at least for catalogue-genes included in the analysis, there is a highly positive correlation in co-expressions between the different tissue-GCNs (Figure 2 & 5). Moreover, leaf and berry GCNs diverge one from another in terms of the co-expressions that are able to capture (Figure 2). This means that aggregated tissue-specific GCNs can be used as a proxy of tissue-specific regulatory networks, since they are able to detect tissue-specific co-expressions. On the other hand, TI GCNs capture co-expressions that are present across all analyzed tissues. This occurs because berry and leaf are the tissues that are contributing the most to the TI GCN (67 out of 131 and 42 out of 131 SRA studies for berry and leaf respectively). Therefore, TI GCN is a suitable tool for analyzing general regulatory networks. For example, the berry GCN captures more efficiently than the leaf GCN the co-expressions of the 5.9 MapMan bin, related to lipid metabolism (Sup. Figure 4). Lipid metabolism is expected to be more associated with berry because changes in lipid metabolism can affect the accumulation of secondary metabolites such as anthocyanins (Duan et al., 2019). On the contrary, leaf GCN performs better on the 27.1 MapMan bin, related to the circadian clock system. This is explained because daily variations of the ripening berry transcriptome is only partially controlled by circadian clock components, being also affected by environmental changes (Carbonell-Bejerano et al., 2014). In these two examples, TI GCN performance is in between the leaf and berry networks.

### GCNs as a proxy of gene regulatory networks

Even if percentage-transformed values can be used to compare GCNs, the best way to compare them between the different tissue datasets is by inspecting the ontology terms enriched in the list of co-expressed genes (as in Figure 4 compared to Sup. Figure 5 and Sup. Figure 6). Indeed, enrichment analysis on a TF gene-centered network is a powerful tool for predicting regulatory relationships. By applying this methodology to all the members of TF gene families, we were able to recover most of the known regulatory relationships for R2R3-MYB and bHLH(MYC) genes, independently of the tissue (Figure 4 & Sup. Figure 5). For example, *MYB14* and *MYB15* (Höll et al., 2013; Orduña et al., 2022) were ascertained to be involved in stilbenoid and p-coumaroyl-CoA biosynthesis, *MYBA1/A2/A6* (Kobayashi et al., 2004; Matus et al., 2017; Walker et al., 2007) were identified as potential regulators of anthocyanin biosynthesis, *MYBPA1/PA2* (Bogs et al., 2007; Terrier et al., 2009) appeared to be involved in the biosynthesis of proanthocyanidins, and *MYC1* (Hichri et al., 2010) showed a relation to the flavonoid biosynthesis pathway. All these GCN-predicted roles were correctly assigned when compared to the literature. For the case of *MYBA7*, an additional known-anthocyanin regulator, only the leaf and TI networks corroborated its role. The berry GCN could not as expected since MYBA7 is a regulator of pigment accumulation only in vegetative organs (Matus et al., 2017). Overall, our findings prove our methodology to be robust enough for detecting true biological regulatory networks. Besides already described regulatory relationships, our methodology was also able to predict novel regulatory relationships.

### *MYBPA1* is a regulator of p-coumaroyl-CoA and proanthocyanidin biosynthesis

The combined use of GCNs and DAP-seq data allowed us to suggest that MYBPA1 could be involved in the increase of the metabolic flux towards the proanthocyanidin biosynthesis starting at the regulation of the shikimate pathway. The initial exploration of the GCNs already supports this observation. Enrichment analyses of MYBPA1 berry, leaf and TI gene-centered networks pointed out involvement in all the steps of the proanthocyanidin biosynthesis, completely matching previously available literature (Bogs et al., 2007; Terrier et al., 2009). In addition, our data suggest MYBPA1 as a potential regulator of p-coumaroyl-CoA biosynthesis at the early steps of the phenylpropanoid pathway. This is a new regulatory relationship that has not been previously tested. MYBPA1 3D plot (Figure 5) confirms the enrichment analysis results, as several genes involved in p-coumaroyl-CoA (*DAHPS3*, *SK5*, *PAL1-2, PAL5*, *C4H1*, *4CL8* and *4CL12*) and proanthocyanidin (*CHS1-2-3*, *CHI1*, *F3H2*, *F3H5*, *F3’H1*-*2*, *CytoB5*, *DFR1*, *LDOX1*, *ANR1* and *LAR1*) biosynthesis are shared between the TI, berry and leaf MYBPA gene-centered networks with high co-occurrence values. In addition, MYC1, whose direct interaction with MYBPA1 has been described (Hichri et al., 2010) appears in the MYBPA1 berry and TI gene-centered networks.

MYBPA1 DAP-Seq results validate the gene-centered networks. Firstly, the MYBPA1 binding motif generated by analyzing the top 600 most significant peak sequences (Figure 6 A) is very similar to the MYB14 and MYB15 binding motifs (Sup. Figure 7), that were generated using the same methodology (Orduña et al., 2022). This similarity may explain why MYBPA1 targets the same genes of the shikimate and early phenylpropanoid pathway as MYB14/15. On the other hand, small differences in the binding motif and bigger divergence in the C-terminal end of MYBPA1 and MYB14/15 may explain the specificity for the regulation of proanthocyanidin and stilbenoid biosynthesis, respectively. Secondly, at least one isoform responsible for each metabolic step of the shikimate and early phenylpropanoid pathways is bound by MYBPA1 (Figure 6 D). This supports the idea that MYBPA1 could be binding and potentially regulating genes from the shikimate and early phenylpropanoid pathway. The robustness of MYBPA1 target identification is supported by the fact that all MYBPA1-characterized targets related to the proanthocyanidin biosynthesis (Bogs et al., 2007; Hichri et al., 2010; Koyama et al., 2014; Terrier et al., 2009) are bound by and co-expressed with MYBPA1 (Figure 6 C). It is also remarkable that DAP-Seq results show a binding peak in the promoter region of the *MYBPA1* gene, suggesting a *MYBPA1* autoregulatory loop. As the GCNs cannot be used for analyzing autoregulation, this mechanism should be further validated experimentally.

The overlap between MYBPA1 gene-centered networks and DAP-Seq dataset in the form of high confidence targets (HCTs) shows a great overlap of genes involved in the shikimate (*DAHPS3-4*, *ESPSP4*, *CS3*, *CM3*, *PA1* and *AD2-3-4*), early phenylpropanoid (*PAL1-2*, *PAL5*, *C4H1*, *4CL1*, *4CL8* and *4CL12*) and proanthocyanidin (*CHS1-2-3*, *CHI1*, *CHI4*, *F3H2*, *F3H5*, *CytoB5*, *DFR1*, *LDOX1*, *ANR1* and *LAR1*) biosynthesis (Figure 6 C & D). Interestingly, *LAR1* appears within the HCTs involved in the PA biosynthesis across the three GCNs. Previous studies suggested *LAR1* as exclusively regulated by MYBPA2, whereas *LAR2* was described to be regulated only by MYPAR and MYBPA1 (Koyama et al., 2014). On the contrary, our results suggest that MYBPA1 is able to bind and upregulate both *LAR* paralogues. Indeed, *LAR2* is co-expressed with MYBPA1, having the same co-occurrence value as the last gene of the co-expression list (i.e., in the position 420), but falling out of this list. In addition, the CHS gene family is detected as HCT (*CHS1*) and VHCT (*CHS2/3*). The fact that *CHS1* is not a VHCT can be explained by the absence of a probe for this gene in the microarray chip. Remarkably, each *CHS* has a different DAP-Seq binding profile (Figure 6F), which corroborates previous literature suggesting different levels of control towards these three paralogues (Harris et al., 2013). As seen in the GCN and the DAP-seq analysis, there is a total absence of genes of the *stilbene synthase* (*STS*) family and also key genes for anthocyanin biosynthesis, such as *UFGT1*. Altogether, there results also imply that the MYBPA1-mediated metabolic flux is favored at a sub-branch level, reducing the accumulation of resveratrol or anthocyanins, which could deviate the metabolic flux away from proanthocyanidins.

Several transcriptional regulators involved in PA biosynthesis appeared to be present among the HCTs. For instance, *MYC1* being an HCT for berry and TI-analysis supports previous studies that showed that *MYC1* promoter activation is dependent on the co-expression of both *MYC1* and *MYBPA1* (Hichri et al., 2010). Previous studies showed that *MYBPA2* overexpression results in *MYBPA1* accumulation, and despite not being able to prove that *MYBPA1* could directly modulate *MYBPA2*, they suggested that *MYBPA2* was acting upstream of *MYBPA1* (Terrier et al., 2009). Here we suggest, conversely, that *MYBPA1* is able to modulate *MYBPA2* expression, placing both TFs at the same hierarchical level. In the same way, *MYBPAR*, also characterized as a PA biosynthesis regulator acting upstream of *MYBPA1* (Koyama et al., 2014), is shown here as a HCT across the three GCNs, suggesting that MYBPA1 is also able to regulate its expression. Thus, our data reevaluate previously established TF-TF hierarchies, however, further experimental validation is still necessary.

The use of *MYBPA1*-overexpression microarray data further allowed the classification of ‘very high confidence targets’ (VHCTs) from a subset of identified HCTs. Here, we give more strength to the hypothesis that MYBPA1 is increasing the metabolic flux towards proanthocyanidin biosynthesis, not only by upregulating genes of the flavonoid pathway involved in proanthocyanidin biosynthesis, but also by upregulating genes from the shikimate and early phenylpropanoid pathways, potentially leading to an increase of *p*- coumaroyl-CoA precursor (Figure 6 C & D). VHCTs are most likely underestimated as some key enzymatic and regulatory genes of the examined pathways are not represented in the microarray chip.

*MYBPA1* validation is a great proof of concept of the potential of the methodology developed in this work for using GCNs as prediction tools of TFs potentially involved in secondary metabolism. As also shown in previously published works (D’Incà et al., 2021; Orduña et al., 2022; Zhang et al., 2021), gene-centered networks extracted from our generated GCNs can be used as validation tools for TFs regulatory roles. The overlap between cistromic datasets (i.e., DAP-Seq), gene-centered networks and transcriptomic datasets (i.e., microarray or RNA-Seq extracted from overexpression experiments) presented here constitutes a robust methodology for the characterization of novel as well as partially established TF regulatory roles.

### aggGCN: An online and interactive tool for exploring GCNs

Due to the utility of our GCNs in serving as a proxy of biological regulatory networks, we developed a user-friendly online resource that allows grape (or any other plant) researchers to publicly access and navigate through all GCNs generated in this study. The Rshiny-based application program (App) named aggGCN, located within the Vitviz platform (http://www.vitviz.tomsbiolab.com), ensures the Findability, Accessibility, Interoperability and Reusability (FAIRness) of the data generated in this study. Different features make aggGCN the best available tool for exploring co-expression relationships: it is interactive, dynamic, and all the data is easily downloadable. The interactive networks tab from our aggGCN app can be used as a useful exploratory tool of pathway-centered networks. This tab allows a visual but biologically meaningful way of subsetting network data, allowing to predict novel regulators of a given metabolic route. The individual GCN tab can be used for creating and downloading gene-centered networks for any PN40024 gene. In the near future, other tissues besides leaf and berry, and cultivar-specific GCNs (especially for berries) will be eventually generated and incorporated to the app when the number of publicly available datasets is sufficient to permit GCN construction. It is worth highlighting that the GCNs described in this study are rapidly updatable with the addition of new SRA studies and can be adapted on new genome assemblies and annotations. AggGCN will be annually updated to integrate newly published RNA-Seq datasets. Finally, we also offer the pipeline developed for GCNs construction to the grapevine and other plant communities, since this pipeline is easy to adapt to any other species of interest if enough transcriptomic datasets are available.

## SUPPLEMENTARY DATA

**Sup. Figure 1**. Workflow of the pipelines for A) downloading, trimming, aligning and filtering of SRA runs and B) for generating aggregated and single GCNs.

**Sup. Figure 2**. Workflow for the analysis of dataset size impact on the network performance. The initial subset to build and evaluate GCNs consisted of two randomly selected SRA studies after which further SRA study subsets were randomly added, generating subsets of the following sizes: TI GCN (2, 5, and 10 increments for ranges 2 – 10, 15 – 50, 60 – 120, respectively), berry GCN (increments of 2 and 5 for ranges as above), and leaf GCN (2 and 4 increments for ranges 2 – 10 and 14 – 38, respectively). For each subset evaluated, this process was repeated 5 times. Additionally, the full set of SRA studies used for a particular GCN (i.e. 131, 67, and 42 for TI, berry, and leaf dataset, respectively) is also included

**Sup. Figure 3**. AUROC of the generated GCNs across each individual GO ontology.

**Sup. Figure 4**. Scatterplot of MapMan bins z-scores across berry, leaf and TI GCNs.

**Sup. Figure 5. MYB gene family Secondary metabolism networks.** A) MYB family berry secondary metabolism network. B) MYB family leaf secondary metabolism network. Squares depict MYB genes, whereas circles depict MapMan bins involved in the secondary metabolism. Circle sizes are determined by the number of MYB genes whose gene-centered networks are enriched in the respective MapMan bin. Red edges depict regulatory relationships already described in the literature.

**Sup. Figure 6. MYC/bHLH gene family Secondary metabolism networks.** A) bHLH family leaf secondary metabolism network. B) bHLH family berry secondary metabolism network. C) bHLH family TI secondary metabolism network. Squares depict MYB genes, whereas circles depict MapMan bins involved in the secondary metabolism. Circle sizes are determined by the number of MYB genes whose gene-centered networks are enriched in the respective MapMan bin. Red edges depict regulatory relationships already described in the literature.

**Sup. Figure 7**. Motif similarity comparison between MYBPA1 and MYB14/15.

**Sup. Table 1**. Table of AUROC values for the generated GCNs across each MapMan bin and GO term used for GCNs evaluation.

**Sup. Table 2**. Table of AUROC values for each individual SRA study single GCN used in the construction of the berry, TI and leaf GCNs.

**Sup. Table 3**. MYBPA1 DAP-Seq peaks, MYBPA1 berry gene-centered network, MYBPA1 leaf gene-centered network, MYBPA1 TI gene-centered network, Terrier et al., 2009 differentially upregulated genes, High Confidence Targets list & Very High Confidence Targets lists.

**Sup. Table 4**. Shared and unique significant relationships between MYB genes and secondary metabolism bins across the berry, leaf and TI GCNs.

**Sup. Dataset 1.** Ontologies used for AUROC metric evaluation in EGAD R package format.

**Sup. Dataset 2.** Manually curated MapMan ontology used for performing enrichment Analysis.

**Sup. Dataset 3.** Shikimate and phenylpropanoid pathway plots in html format.

## KEY WORDS & ABBREVIATIONS

GCN: Gene co-expression network. This term describes co-expression networks built at genomic scale. Therefore, they contain co-expression information between all the genes described in the genome.
Aggregated network: GCN built using network co-occurrence aggregation. Briefly, this aggregation method consists on generating one network per SRA study, and counting across how many SRA studies the same co-expressions are detected.
Single network: GCN built using data aggregation. Briefly, this aggregation method consists on merging the information contained on each SRA study in a unique count matrix and generating a network from the merge matrix count.
Gene-centered network: List of the 420 genes (roughly 1% of the genes described in the PN40024 assembly) being most strongly co-expressed with a gene of interest.
Pathway-centered network: Graphical representation of co-expressions between genes involved in a metabolic pathway and any other gene, including the transcription factors potentially involved in the regulation of that pathway.
Interaction plot: Interactive representation of a pathway-centered network.

## ACKNOWLEDGMENTS

The generation of the networks and all the bioinformatic analysis were performed on the HPC cluster Garnatxa at Institute for Integrative Systems Biology (I2SysBio).

## AUTHOR CONTRIBUTIONS

TM and DW designed the research; LO, DN and AS conducted all the code writing and bioinformatic analysis; CZ performed the DNA Affinity Purification Sequencing experiment; LO, DN, TM and DW wrote the paper. All authors contributed to interpretations and revisions of the manuscript.

## DATA AND CODE AVAILABILITY

The code used for downloading, trimming, aligning and counting of raw counts is publicly available at https://github.com/Tomsbiolab/raw_counts_generation. The code used for generating aggregated GCNs is available at https://github.com/Tomsbiolab/agg_WGCN. The code used for generating single GCNs is available at https://github.com/Tomsbiolab/non_agg_WGCN. The generated apps are available at the Vitviz platform (http://vitviz.tomsbiolab.com/). In addition, AggGCN will become available at the Grape Genomics Encyclopedia (GRAPEDIA; https://grapedia.org/) where it will be used along with different application program interfaces including reference genome browsers, gene cards, transcriptomic data visualizations and software for ‘variation-gene expression-phenotype’ associations.

DAP-Seq data can be found in the Gene Expression Omnibus (GEO) database of the NCBI under the accession GSE230186. All relevant data can be found within the manuscript and its supporting materials.

