## Supplementary figures and images for "Aggregated gene co-expression networks for predicting transcription factor regulatory landscapes in a non-model plant species"

### Sup. Figure 1

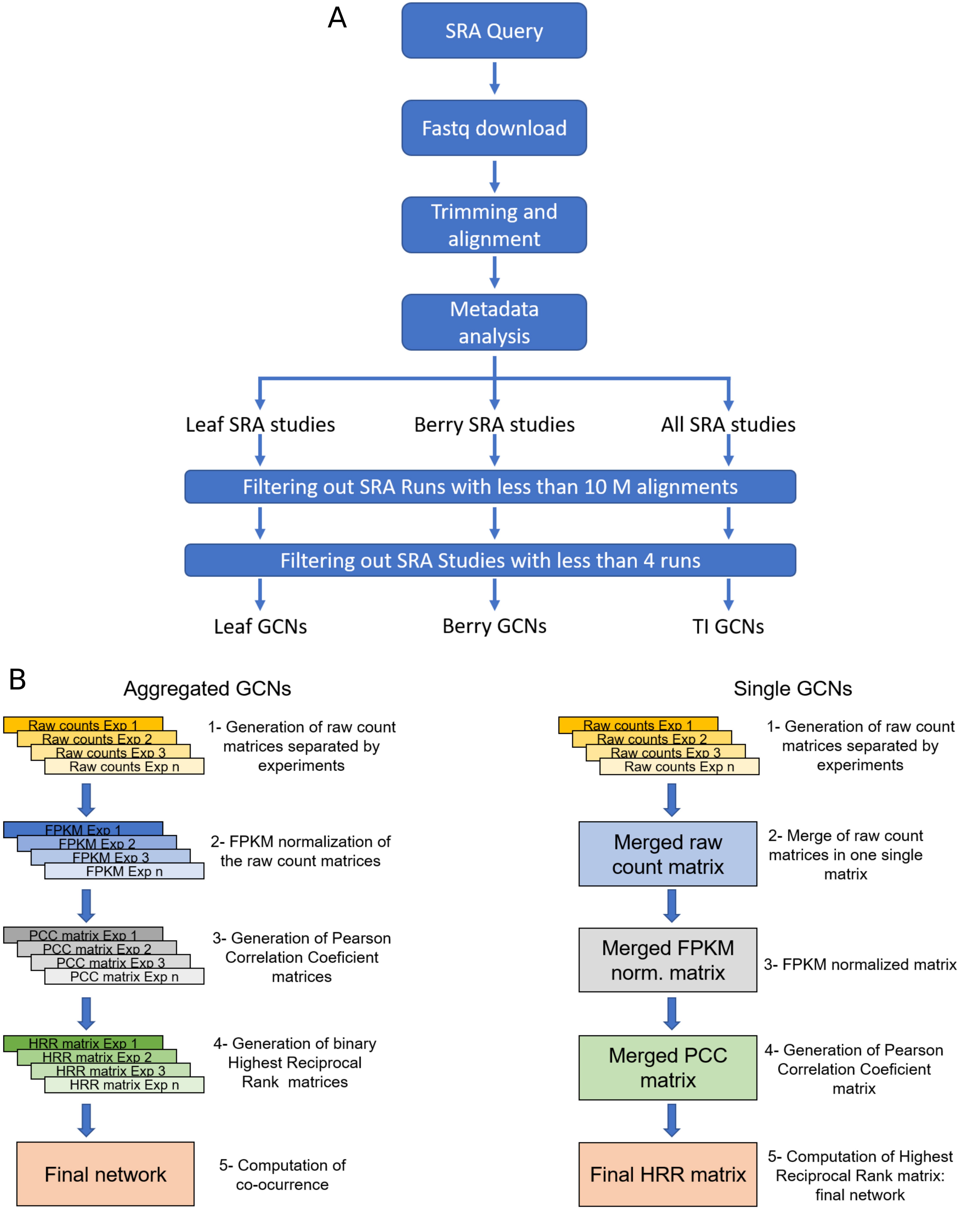

### Sup. Figure 2

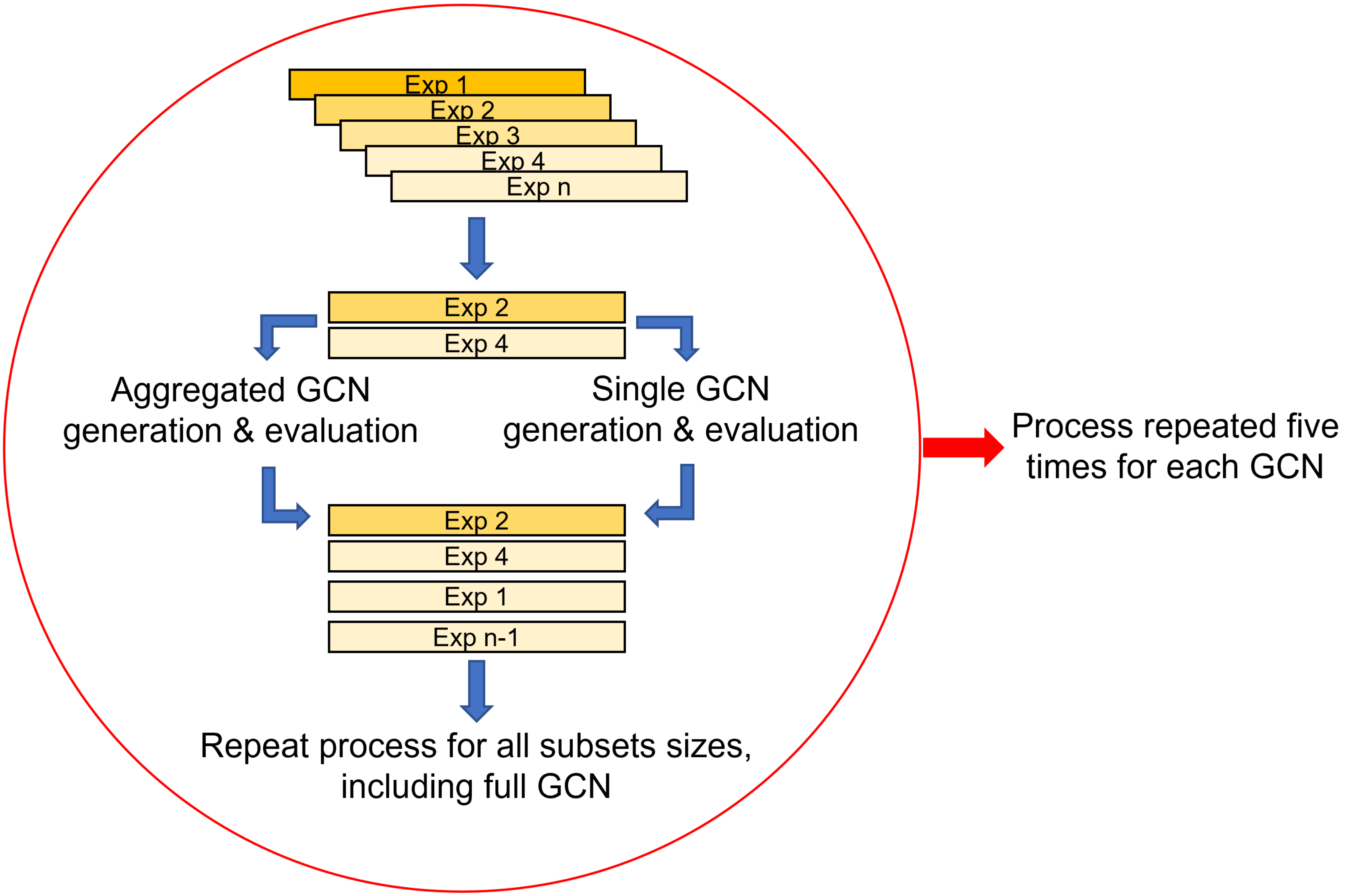

### Sup. Figure 3

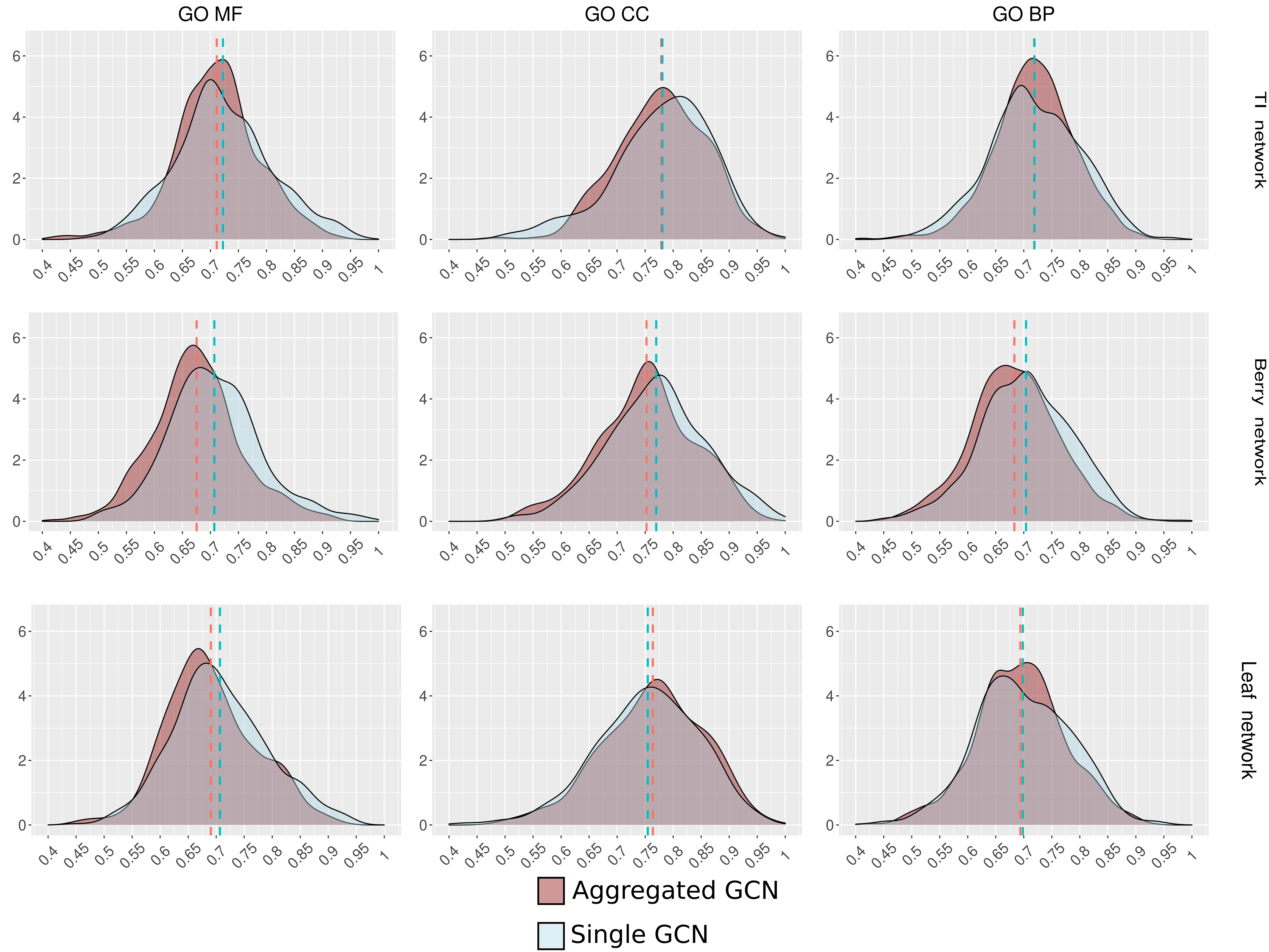

### Sup. Figure 4

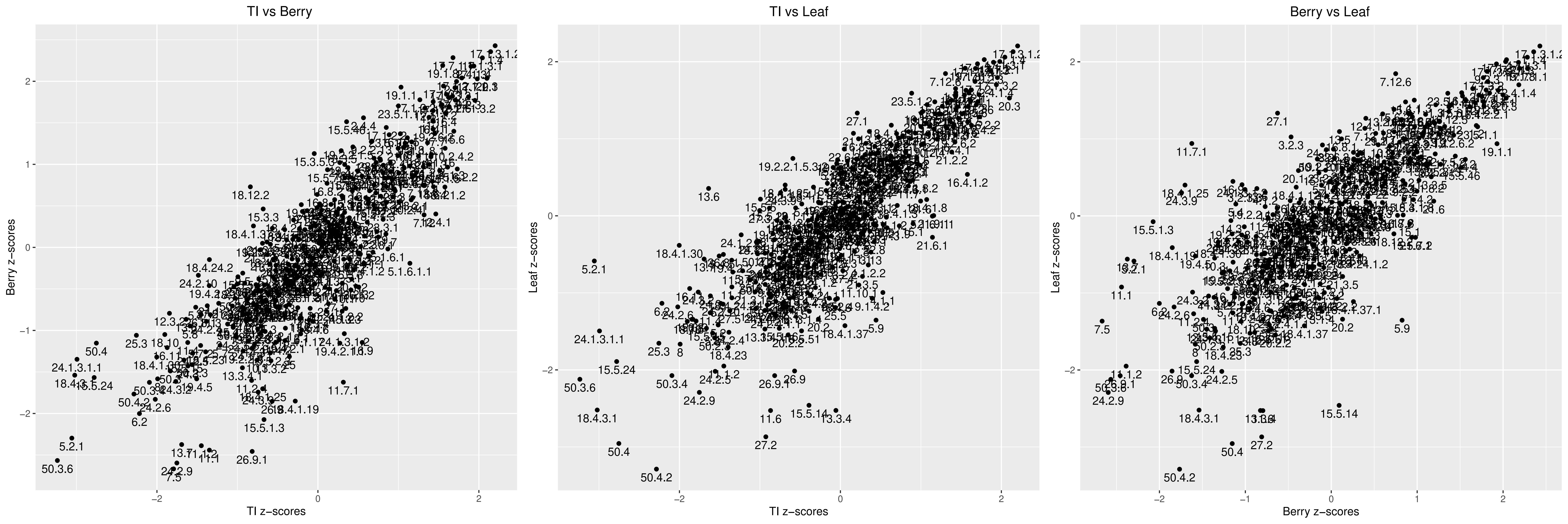

### Sup. Figure 5

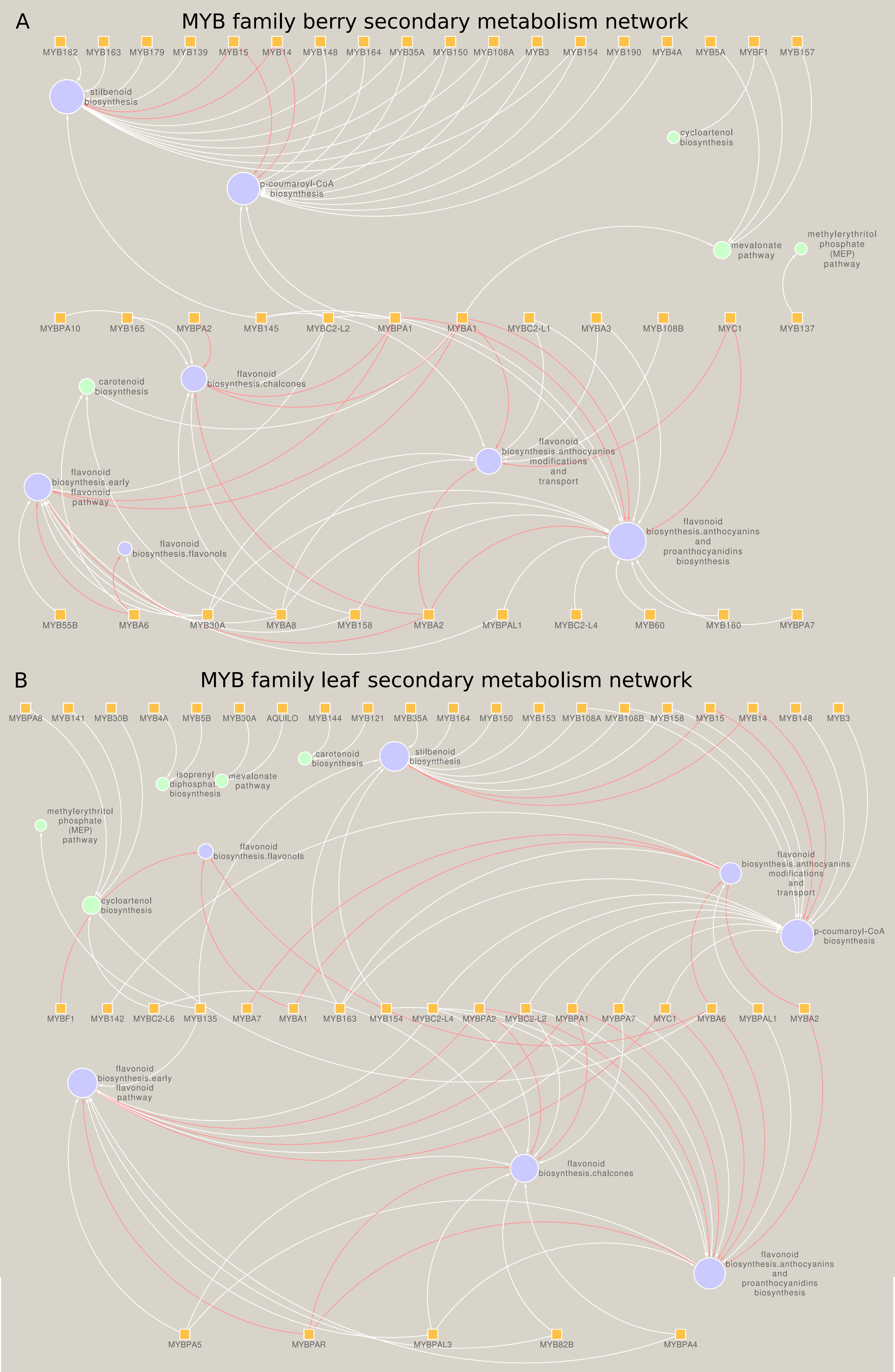

### Sup. Figure 6

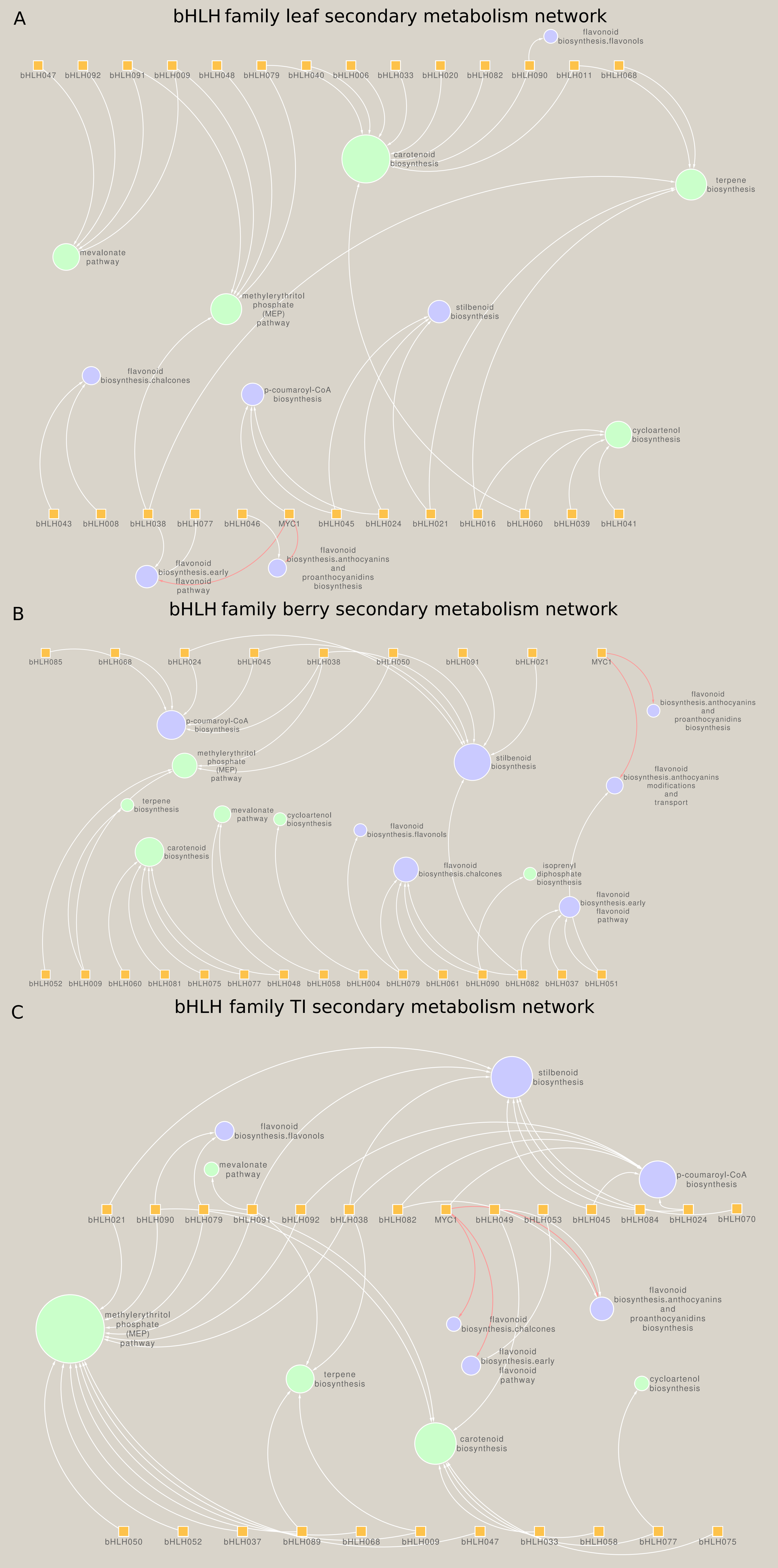

### Sup. Figure 7

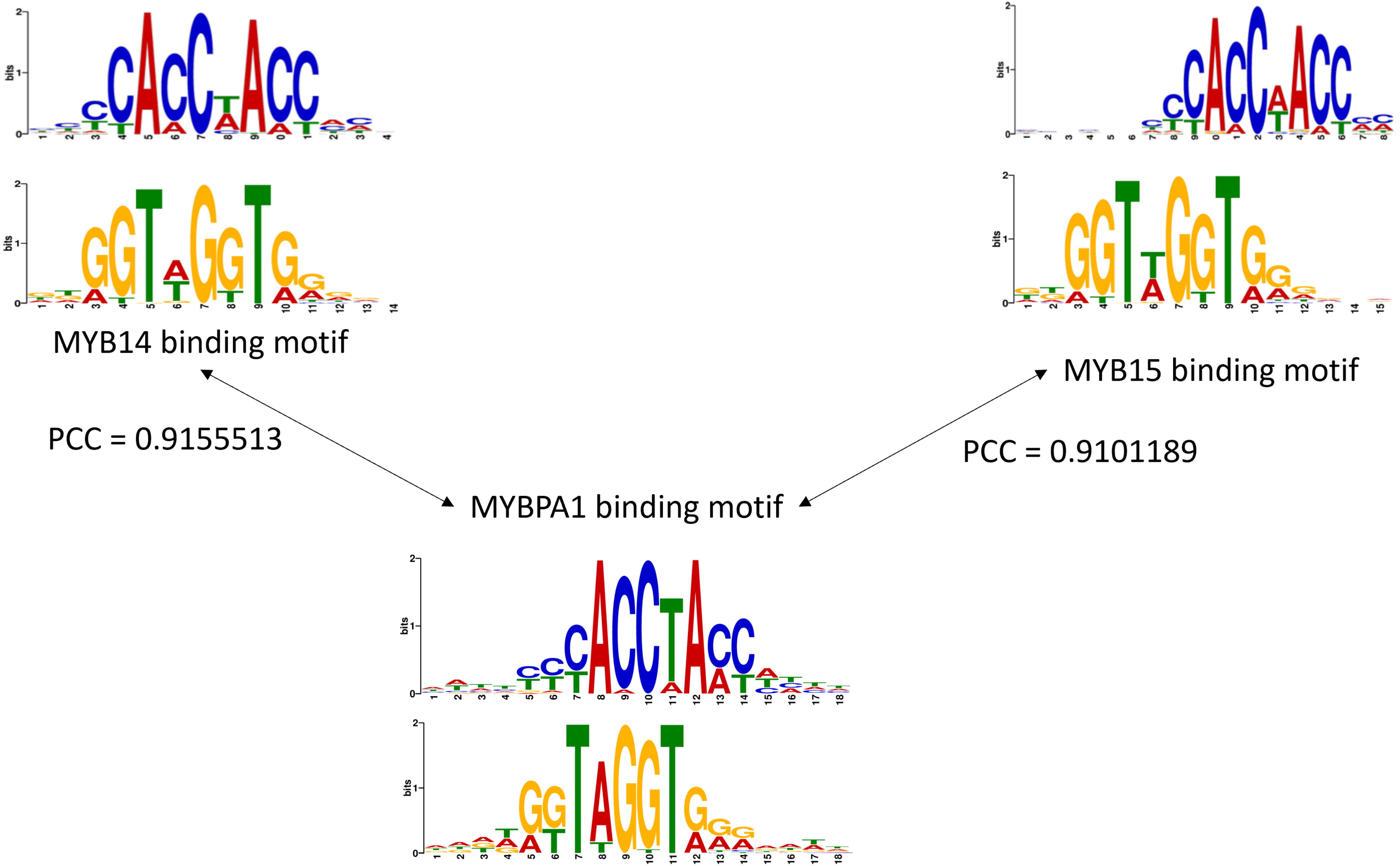
